# Enhancer-mediated metabolic pre-patterning defines trabecular cardiomyocyte identity prior to morphogenesis

**DOI:** 10.64898/2026.02.12.705356

**Authors:** Costantino Parisi, Shikha Vashisht, Mohammad Salar Ghasemi Nasab, Kandhadayar Gopalan Srinivasan, Katarzyna Misztal, Marcin Zagorski, Cecilia Winata

## Abstract

Cardiac trabeculae are essential structures of the vertebrate heart critical for ventricular wall integrity and contractility. Despite their importance, the transcriptional regulatory program defining trabecular cardiomyocyte identity and driving their development remains poorly understood. We identify a novel enhancer, *-6.8got2b*, that drives gene expression specifically in the trabecular myocardium of the zebrafish heart. Functional analyses revealed that *- 6.8got2b* is required for trabeculae formation, likely acting through its target gene, *got2b*, which encodes a mitochondrial enzyme. We delineated a 166bp core region responsible for restricting gene expression to trabecular cardiomyocytes, harboring binding motifs for multiple transcription factors (TFs), including Tbx20, whose motif deletion abolished trabeculae-specific expression. A human genomic region sharing partial sequence similarity and TF motifs with *–6.8got2b* was found near GOT2P4 – a pseudogene related to the human GOT2 gene. Strikingly, the human sequence drove expression in the zebrafish trabeculae and ventricular myocardium, indicating partial but functionally relevant conservation. Using *– 6.8got2b* as a molecular marker, we transcriptionally profiled trabecular cardiomyocytes and show that, compared to compact layer cardiomyocytes, they exhibit distinct gene signatures indicative of transcriptional priming for elevated energy metabolism prior to the formation of overt trabecular structure. Our findings reveal a conserved –*6.8got2b-got2b* regulatory axis driving trabecular cardiomyocyte development downstream of Tbx20, establishing a new layer of transcriptional regulation in heart morphogenesis.

## Introduction

An evolutionarily conserved gene regulatory network is responsible for the intricate morphogenetic steps required for the formation of the heart. A core collection of conserved cardiac transcription factors (TFs) is involved in heart cell fate determination, contractility, morphogenesis, segmentation, and growth^1,2^. These TFs govern and stabilize the heart’s developmental program^3–5^. Despite the wealth of information on TFs involved heart development, the regulatory elements through which they exert their functions remain poorly defined. Large-scale projects such as ENCODE^6^, the Pediatric Cardiac Genomics Consortium^7^, Pediatric Heart Network^8^ and the UK10K consortium^9^ have identified numerous putative regulatory elements involved in cardiac development and disease. However, only a few of these elements have been functionally validated or studied in detail to understand their functional mechanism.

Enhancers are *cis* regulatory elements in the genome that regulate gene expression in a precise and dynamic manner. They can be located at any distance from their target genes and make physical contact with gene promoters through chromatin looping, enabling them to directly influence transcription output^10–12^. TFs bind to enhancers and mediate gene regulation by recruiting other proteins and enzymes that modify chromatin structure and initiate transcriptional activation^13,14^. Unlike gene promoters, enhancers lack distinctive sequence motif and fixed genomic location and are thus much more difficult to identify. Various approaches have been developed to identify enhancers at genome-wide scale using several defining criteria. These include the presence of enhancer-associated histone modification marks such as H3K4me1 and H3K27ac^15,16^, open chromatin regions, as well as the binding of TFs^17,18^.

While evolutionary sequence conservation within the non-coding region has been commonly used to indicate developmental enhancers^19^, recent studies increasingly show that many functional enhancers are resistant to extensive sequence divergence while still retaining their tissue-specific activity and function^20,21^. Instead of strict sequence conservation, these enhancers often preserve combinatorial TF binding sites, implying the preservation of protein-protein interactions that sustain their collective occupancy^22–24^. Notably, functional analyses of orthologous enhancers across species reveal that many enhancers initially identified for driving specific gene expression in mammals retain similar activity in fish, highlighting conserved regulatory mechanisms across evolutionary lineages^25–28^.

Cardiac trabeculae are an essential feature of the vertebrate heart which play critical roles in oxygen and nutrient uptake in the embryonic heart, ventricular contraction, and overall heart health^29^. Improper trabeculae development can result in impaired cardiac growth, leading to embryonic lethality or congenital heart disease. Disruption of the balance between proliferation in the compact and trabecular layers of the ventricular wall can cause cardiomyocyte hyperplasia, manifesting as hypoplastic left heart syndrome, which accounts for up to 3% of all congenital heart disease cases in infants^30^. On the other hand, excessive trabeculation is linked to certain forms of cardiomyopathy, including left ventricular noncompaction cardiomyopathy (LVNC)^31^.

The formation of cardiac trabeculae begins during embryonic development when the heart becomes functionally active. They emerge as an extension of the ventricular myocardium, forming a sponge structure in the embryonic heart. As the heart grows, the muscular walls of the ventricles undergo extensive remodelling, where the trabecular structure is compacted, marking the maturation of the heart^32–34^. Trabeculae development relies on a complex series of signalling events involving multiple developmental pathways and includes cellular crosstalk between the endocardium, myocardium, and cardiac extracellular matrix (reviewed Qu et al., 2022^35^). During trabeculae development, cardiomyocytes undergo extensive changes, which require energy metabolism^36^, involving the rapid production of ATP and other intermediates necessary to meet the high-energy demands of cell proliferation and migration in developing tissues^37^.

The mechanisms and the signalling crosstalk underlying cardiac trabeculae formation in mammals have been shown to be conserved in the zebrafish^38–40^, making it an excellent model to study this process *in vivo*. In the zebrafish heart, trabeculae formation is initiated at around 60 hpf, when CMs in the outer curvature of the ventricle delaminate, migrate from the compact layer, and proliferate, forming multicellular sponge-like projections^40–43^. Multiple signalling pathways between the myocardium and endocardium have been implicated in regulating cardiac trabeculation, including Nrg2a/Erbb2^39,44^, Notch^45^, Hippo^46^ and Apelin^47^, as well as physical forces generated by cardiac contractility and blood flow^45,48^. Despite these advances, the upstream regulatory mechanism that initiate the earliest changes in trabecular cardiomyocyte behaviour and govern their subsequent morphogenesis remain poorly understood. Additionally, the absence of specific markers for these cells further limits the ability to isolate and study them in detail.

Here we identify a novel enhancer, *-6.8got2b*, that drives gene expression specifically in trabecular cardiomyocytes. We show that this enhancer is necessary for cardiac trabeculae formation, and likely acts through its target gene, *got2b*, a gene encoding a mitochondrial enzyme. A core region of the enhancer confers trabeculae-specific expression and requires Tbx20, a transcription factor known for its role in cardiomyocyte development. Comparative genomic analyses revealed a human sequence with partial similarity and shared regulatory features that drove expression in zebrafish trabecular and ventricular myocardium, suggesting functional conservation of the enhancer-gene regulatory axis. Using *–6.8got2b* as a marker, we isolated trabecular cardiomyocytes and defined their early transcriptional landscape. Altogether, our results uncover a conserved *–6.8got2b-got2b* regulatory axis under the control of Tbx20 as a new transcriptional circuit underlying cardiac patterning.

## Results

### Identification of a novel cardiac enhancer *-6.8got2b*

A putative enhancer in zebrafish was identified in our previous study through an integrated analysis of RNA-seq and ATAC-seq data on cardiomyocytes at 24, 48 and 72 hpf embryos^49^. The enhancer, from hereon designated as *-6.8got2b*, is located in chr7:51,767,759-51,768,272 (zebrafish genome assembly GRCz11/danRer11). The *-6.8got2b* enhancer region overlapped with open chromatin regions and the H3K27ac epigenetic mark which indicates active enhancer (**Figure 1A**).

**Figure 1.**
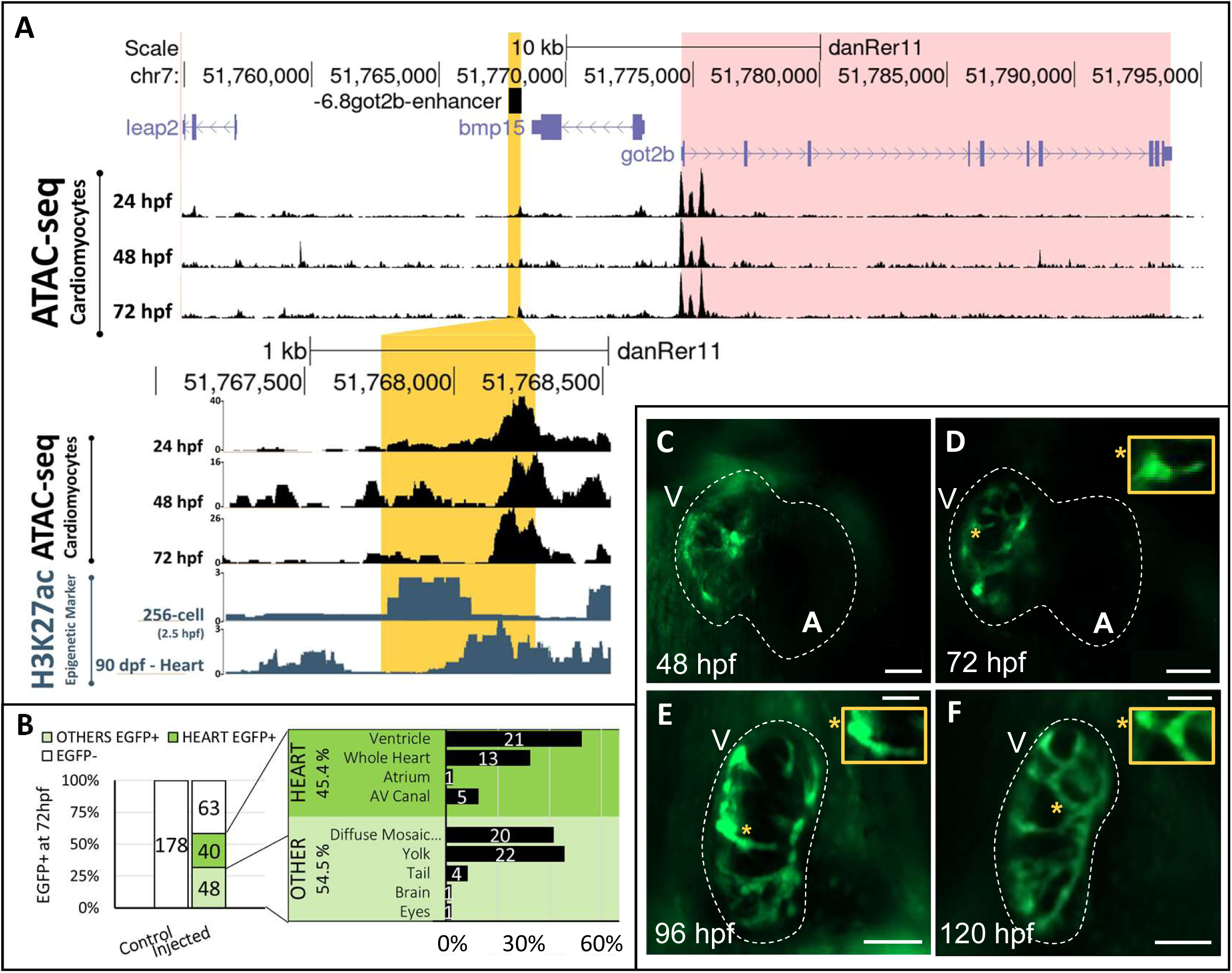
Genomic features and expression pattern driven by the *-6.8got2b* enhancer. A, genome browser view of *-6.8got2b*, with custom tracks for whole cardiomyocytes ATAC-seq data^49^ and H3K27ac chromatin modification marks across various developmental stages. **B**, quantification of reporter expression at various tissues driven by *-6.8got2b* enhancer in transient expression assay; data acquired at 72 hpf from 3 independent experiments. **C-F**, time course imaging reveal EGFP expression in the cardiac ventricle driven by *-6.8got2b* at 48, 72, 96 and 120 hpf. Detailed trabecular morphology is shown in highlighted frame (*); A = Atrium; V = Ventricle; Scale bar = 50 μm.

To test the enhancer activity of *-6.8got2b*, we performed a transient reporter assay using a Tol2-based vector carrying EGFP reporter under the weak *cfos* promoter^50^. A region spanning 514 bp was expected according to the reference genome, however, isolation and sequencing of the region from zebrafish AB and TL wild-type strains revealed a gap of two base pairs (A,T) at position chr7:51767950-51767951 (GRCz11/danRer11), and a variation of A to G at position chr7:51768224-51768224 (GRCz11/danRer11), resulting in a total sequence length of 512 bp (**Supplemental Figure S1**). Thus, a region of 512 bp (chr7:51,767,759-51,768,272; GRCz11/danRer11) comprising the *-6.8got2b* enhancer was cloned into the vector and the resulting *-6.8got2b*-enhancer cfos-EGFP-Tol2 construct was microinjected into zebrafish embryos at the one-cell stage. A total of 473 embryos were injected, of which 151 survived. At 72 hpf, 58.2% (88/151) of injected individuals showed EGFP reporter expression in various domains, among which 45.4% (40/88) primarily exhibited EGFP expression in the heart (**Figure 1B**).

To investigate the spatiotemporal expression pattern driven by the *-6.8got2b* enhancer, stable transgenic lines carrying -*6.8got2b* was generated by raising the 40 embryos exhibiting EGFP+ heart expression, of which 32 survived to adulthood. Five F0 individuals were subsequently outcrossed with wild-type and their offsprings (*F1*) exhibited germ line transmission, consistently expressing the EGFP reporter in the heart. Two stable lines were established from two out of five F0 individuals, and more detailed observation was performed in one line, from hereon designated as Tg(*-6.8got2b-cfos:EGFP)*. Overall, the reporter expression domain in Tg(*-6.8got2b-cfos:EGFP)* encompassed the cardiac ventricle and a restricted domain at the choroid plexus observed at 96 hpf (**Supplemental Figure S2**). Time-course analyses revealed that -*6.8got2b* drove EGFP reporter expression in distinct domains within the cardiac ventricle as early as 48 hpf (**Figure 1C**). By 72 hpf, the EGFP-expressing structure has assumed an elongated morphology (**Figure 1D**, 72 hpf highlighted frame). At 96 hpf, the EGFP+ cells adopted a distinctive finger-shaped appearance (**Figure 1E**, 96 hpf highlighted frame), which interconnected at 120 hpf to form a compact and intricate ridge-like network (**Figure 1F**, 120 hpf highlighted frame).

### The enhancer *-6.8got2b* marks early trabecular cardiomyocytes

To ascertain the identity of the *-6.8got2b*-driven expression domain, the Tg(*-6.8got2b-cfos:EGFP*) line was outcrossed with either Tg(*myl7:mRFP*) which express RFP in the cardiomyocytes (CMs), or Tg(*kdrl*:mCherry) which express mCherry in the endocardium. At 48 hpf, the EGFP-expressing domain overlapped with the RFP signal of the Tg(*myl7:mRFP*) in the monolayered ventricular myocardium (**Figure 2A**, 48 hpf). The myocardial identity of the EGFP-expressing cells was further supported by the clear separation between the *- 6.8got2b*-driven EGFP expression and the endocardial mCherry signal in Tg*(kdrl:mCherry)* (**Supplemental Figure S3**). At 72 hpf, the EGFP-expressing cardiomyocytes appeared more circular and started to protrude out of the compact layer and into the cardiac lumen (**Figure 2A**, 72 hpf, arrows). By 96 hpf, these EGFP-expressing cells occupied the inner ventricular wall, clearly distinguishable from the compact myocardium (**Figure 2A**, 96 hpf), and formed extensive network throughout the ventricle by 120 hpf (**Figure 2A**, 120 hpf, arrows). 3D reconstruction of the Tg(*-6.8got2b:EGFP*) and Tg(*myl7*:RFP) signals (**Figure 2B**) revealed the morphology of the *-6.8got2b*-driven EGFP expression domain within the interior of the ventricular wall which increased in size and complexity over time. At 48 hpf, the structure appeared as distinct subdomains within the ventricular lumen which became more continuous and expanded throughout the length of the ventricle by 72 hpf. At 96 and 120 hpf, this domain appeared as a lamellar structure around the inner ventricular wall (**Figure 2B**). Taken together, our observation suggests that the *-6.8got2b*-enhancer drives expression in a subset of ventricular myocardial cells, which subsequently form organized structures within the inner ventricular wall.

**Figure 2.**
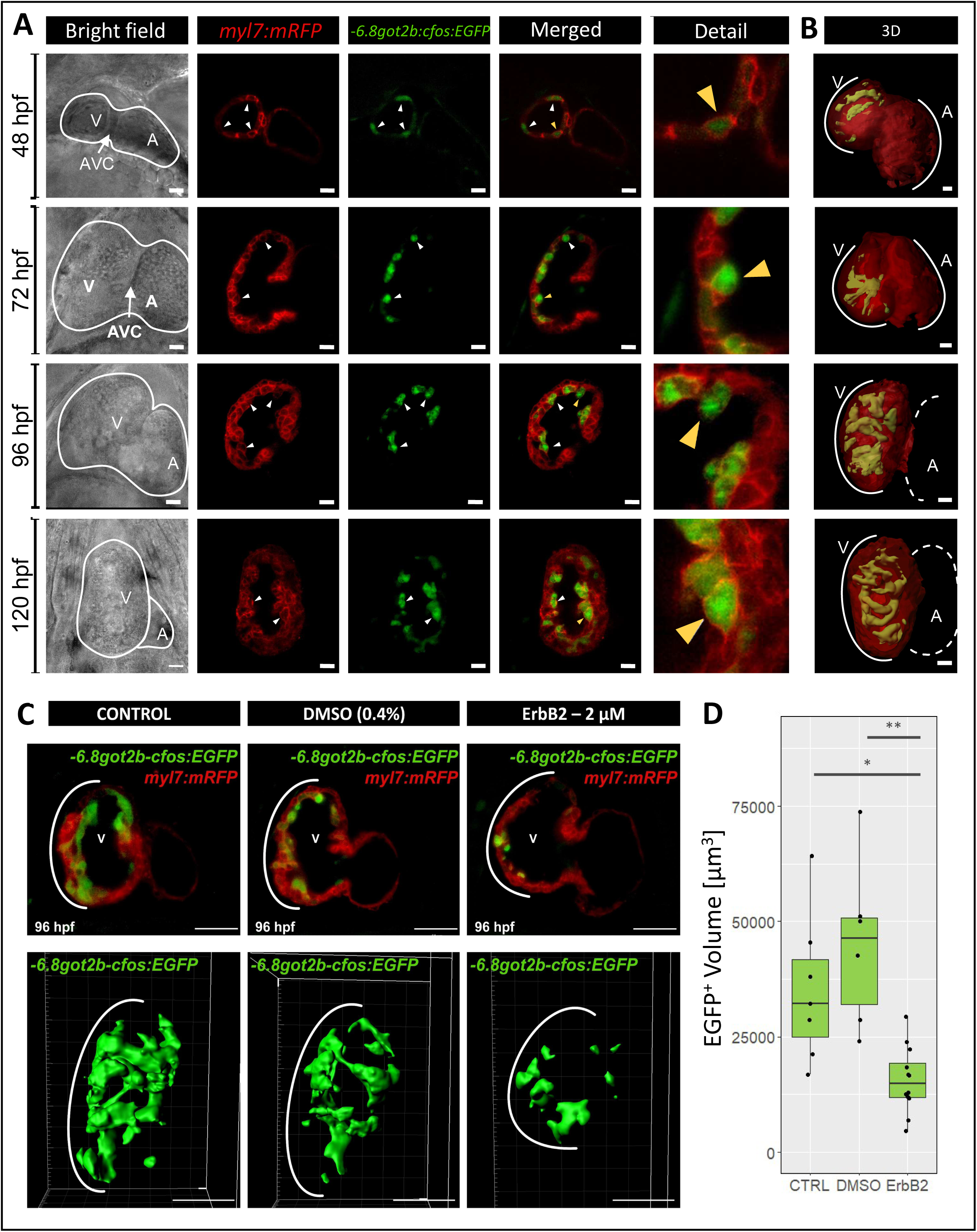
***-6.8got2b* drives expression in trabecular cardiomyocytes. A**, outcross of *Tg(−6.8got2b:EGFP*) with *Tg(myl7:mRFP)* which marks the myocardium at 48, 72, 96 and 120 hpf. A = Atrium; V = Ventricle; AVC = Atrioventricular Canal; Scale bar = 20 μm. **B**, 3D reconstruction of the fluorescent signal from *Tg(–6.8got2b-cfos:EGFP*) x *Tg(myl7:mRFP*) at 48, 72, 96 and 120 hpf; Scale bar = 20 μm. **C**, trabeculae morphology in *Tg(–6.8got2b-cfos:EGFP)* x *Tg(myl7:mRFP)* at 96 hpf, in control, DMSO 0.4%, and inhibitor treatment (2 μM of Erbb2 inhibitor in 0.4% DMSO); V = Ventricle. Below, a representative 3D reconstruction of the EGFP signal for each condition using Imaris software is shown. Scale bar = 50 μm. **D**, comparison of the volume (μm³) of EGFP+ reconstructed structures. Each dot is a replicate (CTRL n = 7, DMSO n = 6, ErbB2 n = 12).; CTRL = control. * P < 0.05; **P < 0.01 Welch’s test.

The localization and morphology of the expression domain driven by *-6.8got2b* resembles that previously described for cardiac trabeculae. To confirm the identity of this domain, we treated the [Tg(*-6.8got2b:EGFP*) x Tg(*Myl7*:mRFP)] larvae with the pharmacological inhibitor AG1478 which temporarily blocks Erbb2 activity and effectively inhibit the development of ventricular trabeculae^36,39^. Exposure of embryos to AG1478 from 96 hpf resulted in a significant reduction of the EGFP-expressing domain in terms of volume and disorganization of its morphology by 120 hpf (**Figure 2C-D**). Inhibitor treatment resulted in low structural complexity and hypomorphic EGFP-expressing structure compared to untreated and DMSO-treated controls. Taken together, the analysis of expression pattern and the reduction of the expression domain by Erbb2 inhibition established that *-6.8got2b* drives specific gene expression in the trabecular cardiomyocytes. Moreover, the reporter expression is observed much earlier than any previous reports of a detectable trabecular structure in the zebrafish heart, positioning *-6.8got2b* activity as a novel early marker of this cell type.

### The enhancer –6.8got2b is necessary for trabeculae development

To investigate the role of the *-6.8got2b* enhancer in trabeculae development, we employed CRISPR-Cas9 based F0 knockout^51,52^ to delete the endogenous enhancer region from the zebrafish genome (**Figure 3A**). The sgRNA-Cas9 complex targeting the enhancer region was microinjected into embryos derived from the outcross of zebrafish lines [Tg*(−6.8got2b:EGFP*) x Tg(*Myl7:mRFP*)] to visualize both the trabeculae and whole myocardium. Out of 188 injected embryos 46.3% (87 out 188) exhibited consistently reduced EGFP expression marking trabecular CMs (**Figure 3B**). A subset of 30 randomly selected embryos from this group were genotyped, revealing that 33.3% (10 out of 30) carried a monoallelic deletion, and 40% (12 out of 30) had a biallelic deletion, resulting in a total of 73.3% of embryos carrying the desired deletion (**Figure 3C**). By 96 hpf, the loss of *-6.8got2b* resulted in a markedly decreased EGFP signal compared to the control siblings, indicating hypomorphic trabeculae. These embryos formed less ridges on their ventricular wall and showed little to no myocardial cell protrusions compared to control conditions, indicating a reduction in overall trabecular structure complexity (**Figure 3D-E**).

**Figure 3.**
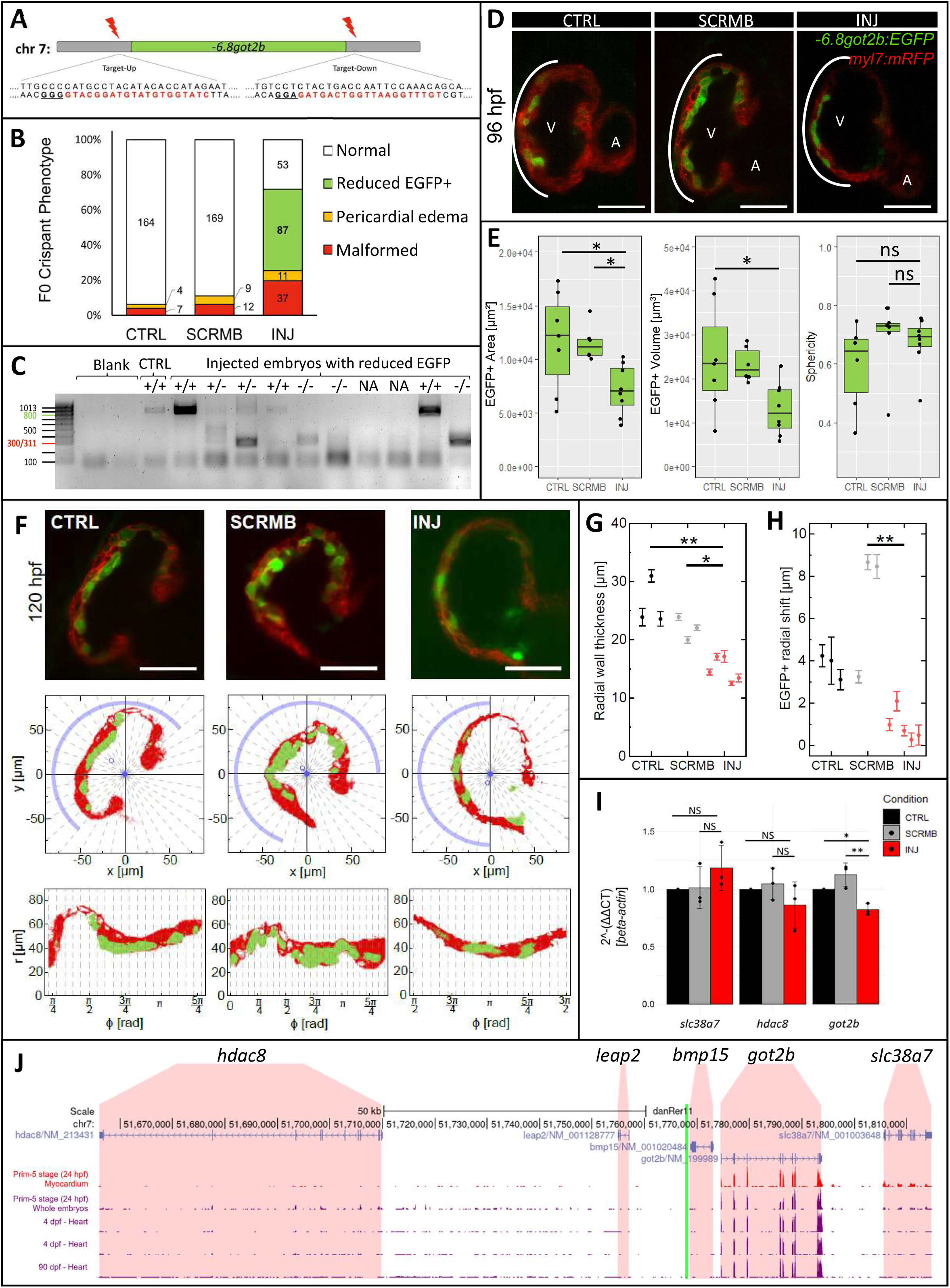
The *-6.8got2b* enhancer regulates cardiac trabeculae development through its downstream target gene *got2b*. **A**, the *–6.8got2b* enhancer region with the gRNA target sequence, with the PAM sequence underlined. **B**, quantification of phenotypes observed at 96 hpf for control (CTRL), as well as embryos injected with scramble (SCRMB) and *s*gRNA targeting *–6.8got2b* region (INJ). **C**, genotyping PCR of representative embryos from the group exhibiting reduced EGFP trabeculation. Expected band size with successful CRISPR deletion is ∼300bp; NA = Not Amplified. **D**, light-sheet image showing cardiac trabeculae morphology of Control, Scramble-Cas9, and sgRNAs-Cas9 groups; Scale bar = 50 μm. **E**, quantification of area (μm²), volume (μm³), and sphericity based on EGFP signal. Each dot represents a replicate; CTRL n = 7, SCRMB n = 6, INJ n = 9. *P < 0.05; one-way ANOVA with Bonferroni’s post hoc test. **F**, representative slices of Z-stack data for CTRL (left), SCRMB (middle) and INJ (right) conditions at 120 hpf, along with 2D spatial analysis. Individual Z-stack slices had background removed and were binarized (mid row). The center of mass (blue circle) was adjusted to mark the center of the heart (blue disk). The latter was used to map original coordinates (x,y) to polar coordinates (r, ϕ). Angular sectors are indicated with dashed lines. Only sectors encompassed by blue arcs were analyzed (bottom row); Scale bar = 50 μm. **G**, radial wall thickness for selected angular sectors for indicated conditions, CTRL (black, n=3, 32 Z-slices), SCRMB (gray, n=3, 50 Z-slices), INJ (red, n=5, 74 Z-slices), mean ± 95% CI. *P < 0.05; **P < 0.01; one-way ANOVA with Bonferroni’s post hoc test. **H**, radial shift of trabecular cells with respect to myocardial cells. Higher values indicated that EGFP+ cells were located more centrally than myocardial cells. Sample sizes and reported significance as in G. **I**, quantitative PCR (qPCR) expression analysis of *slc38a7*, *hdac8*, and *got2b* in response to *–6.8got2b* KO. qPCR was performed on Control, Scramble-Cas9, and sgRNAs-Cas9 groups, with *actb1* used for normalization. Data are presented as mean ± SD (n = 3). *P < 0.05; P < 0.01; one-way ANOVA with Bonferroni’s post hoc test. CTRL = Control, SCRMB = Scramble-Cas9, INJ = sgRNAs-Cas9. **J**, whole cardiomyocytes RNA-seq data^49^ showing expression levels of genes within ±50 kb of the *–6.8got2b* region, transcript levels represented as transcripts per million (y-axis).

To more accurately quantify the changes in myocardial geometry caused by *–6.8got2b* enhancer loss, we developed a computational workflow for image segmentation, spatial analysis, and visualization (Methods). We quantified the spatial distribution of GFP and RFP signals by calculating their centers of mass, providing a measure of their anatomical alignment (**Figure 3F**). Radial wall thickness was estimated by segmenting the heart structure and averaging the measurements of boundary distances within 36 angular sectors per slice for each condition. This approach revealed a significant reduction in wall thickness in *-6.8got2b* loss of function (14.9 ± 0.9 μm; mean ± SEM) compared to controls (24.1 ± 1.5 μm; p = 0.0009, two-sided t-test), indicating impaired myocardial and trabecular development (**Figure 3G**). We then compared the spatial positioning of trabecular and compact myocardial cells by analyzing their average radial distances from the center of the heart. In control embryos, trabecular cells were located significantly closer to the center than myocardial cells, with an average difference of 5.3 ± 1.0 μm (22 ± 4% relative to wall thickness) (**Figure 3H**). In contrast, this separation was markedly reduced in *-6.8got2b* loss of function (0.9 ± 0.3 μm; 6 ± 2%), indicating a disruption in normal trabecular growth (**Figure 3H**). Statistical analysis confirmed a significant difference between crispant and control groups (p = 0.0052, two-sided t-test). To further investigate the spatial distribution of GFP-expressing cells within the myocardium, we determined whether cells increase in size or preferentially shift towards the center of the heart along radial trajectories. We analyzed the spread of GFP and RFP signals within each angular sector. Comparison of the spatial spread between trabecular and myocardial cells showed no significant difference across conditions (p = 0.18) when comparing *-6.8got2b* loss of function to control groups (**Supplemental Figure S4**). This suggests that the observed shift in trabecular position in *-6.8got2b* loss of function is not due to changes in the number of trabecular processes but rather reflects a lack of trabecular structure expansion relative to the compact myocardial layer.

### The *-6.8got2b-got2b* regulatory axis drives trabeculae development

To identify the putative target gene(s) of *-6.8got2b*, we determined whether the expression of any of its neighbouring genes correlated with that driven by the enhancer. The *-6.8got2b* locus falls within a topologically associating domain containing several gene loci within approximately 100 kb region up- and downstream (**Supplemental Figure S5A)**^53^. Directly upstream is *leap2* (−10.6 kb from TSS), followed by *hdac8* (−57.7 kb from TSS). Downstream from this enhancer are *bmp15* (+5.3 kb from TSS), *got2b* (+6.8 kb from TSS) and *slc38a7* (+37.8 kb from TSS). Our previously published transcriptome data^49^ revealed that *hdac8, slc38a7,* and *got2b* were expressed in the developing cardiomyocytes from 24 hpf to 72 hpf, with *got2b* exhibiting a significantly higher expression level across various developmental stages spanning 24 hpf to adult (**Figure 3J).** Furthermore, single-cell RNA-seq analyses of the whole zebrafish heart^54^ revealed the highest expression of *got2b* in cells of the ventricular cardiomyocytes (**Supplemental Figure S5B**). In contrast, *hdac8, slc38a7,* and *leap2* did not exhibit any significant expression in the cardiomyocytes (**Supplemental Figure 5C-E**). In the whole embryo, *got2b* is expressed in several tissues, including the skeletal muscle and cardiovascular system^55^. In case of the latter, its expression was restricted to the cardiac ventricle, which closely resembled that driven by the *-6.8got2b* enhancer. qRT-PCR in the F0 *-6.8got2b* crispants revealed a significant decrease in transcript level of *got2b* compared to uninjected controls (**Figure 3I**). In addition, no significant changes in the expression levels of *hdac8*, *slc38a7,* and *got2a* were observed, supporting the specific regulatory interactions between *-6.8got2b* enhancer and *got2b*. Taken together, these observations supported *got2b* as the likely target gene of the enhancer *-6.8got2b* and suggests their regulatory interaction as a new mechanism underlying trabeculae development.

### Motif analysis reveals core region of the *-6.8got2b* enhancer responsible for specific ventricular trabeculae expression

To determine potential upstream regulators of the *-6.8got2b* enhancer, we searched for TF binding motifs within its sequence. A total of 62 motif occurrences (p < 0.0001) were found within the *-6.8got2b* sequence. Intersection with ATAC-seq data from developing cardiomyocytes^49^ revealed that a large portion of the enhancer region were in an open chromatin conformation in whole cardiomyocytes from 24 hpf through 72 hpf, suggesting its developmental activity (**Table 1**). The TF binding motifs appeared to cluster in three distinct regions of the enhancer, denoted as α, β, and γ (**Figure 4A**). To further pinpoint the minimal region responsible for the *-6.8got2b* enhancer activity, we individually cloned the three distinct regions of the enhancer and tested their activity with the reporter activity assay. The α fragment consisting of 215 bp drove EGFP reporter expression predominantly in the heart (52.4%, 166 out of 314; **Figure 4B**). Of these, 57% (50 out of 87) had expression in the whole heart, while 37.9% (33 out of 87) only in the ventricle. The β fragment of 166 bp drove reporter expression in 69.7% (224 out of 321) injected embryos at 72 hpf, among which, 41.5% (93 out of 224) exhibited expression primarily in the cardiac ventricle (56.9%, 53 out of 93; **Figure 4B**). The second most common expression was in the whole heart (38.7%, 36 out of 93). 3D imaging further revealed that, although the α fragment could also drive expression in the ventricle, its expression domain was in the squamous cells of the compact myocardium, unlike that of the β fragment which was morphologically rounded and localized in the deeper ventricular myocardium characteristic of trabecular CMs (**Figure 4C**). On the other hand, the fragment γ of 178 bp drove predominantly non-specific EGFP reporter expression in only 38.8% (78 out of 201) injected embryos at 72 hpf. Only 20% (15 out of 78) of these exhibited fluorescent expression in the heart, among which, 60% (9 out of 15) showed expression throughout the entire heart, while 40% (6 out of 15) specifically exhibited expression in the ventricle. Intriguingly, although unable to efficiently drive cardiac expression, the γ region overlapped with the highest ATAC-seq signals in whole cardiomyocytes and contained a total of 25 motifs characterised by two dense areas containing TF binding motifs for PPAR and Wnt signalling, respectively (**Figure 4A**).

**Figure 4.**
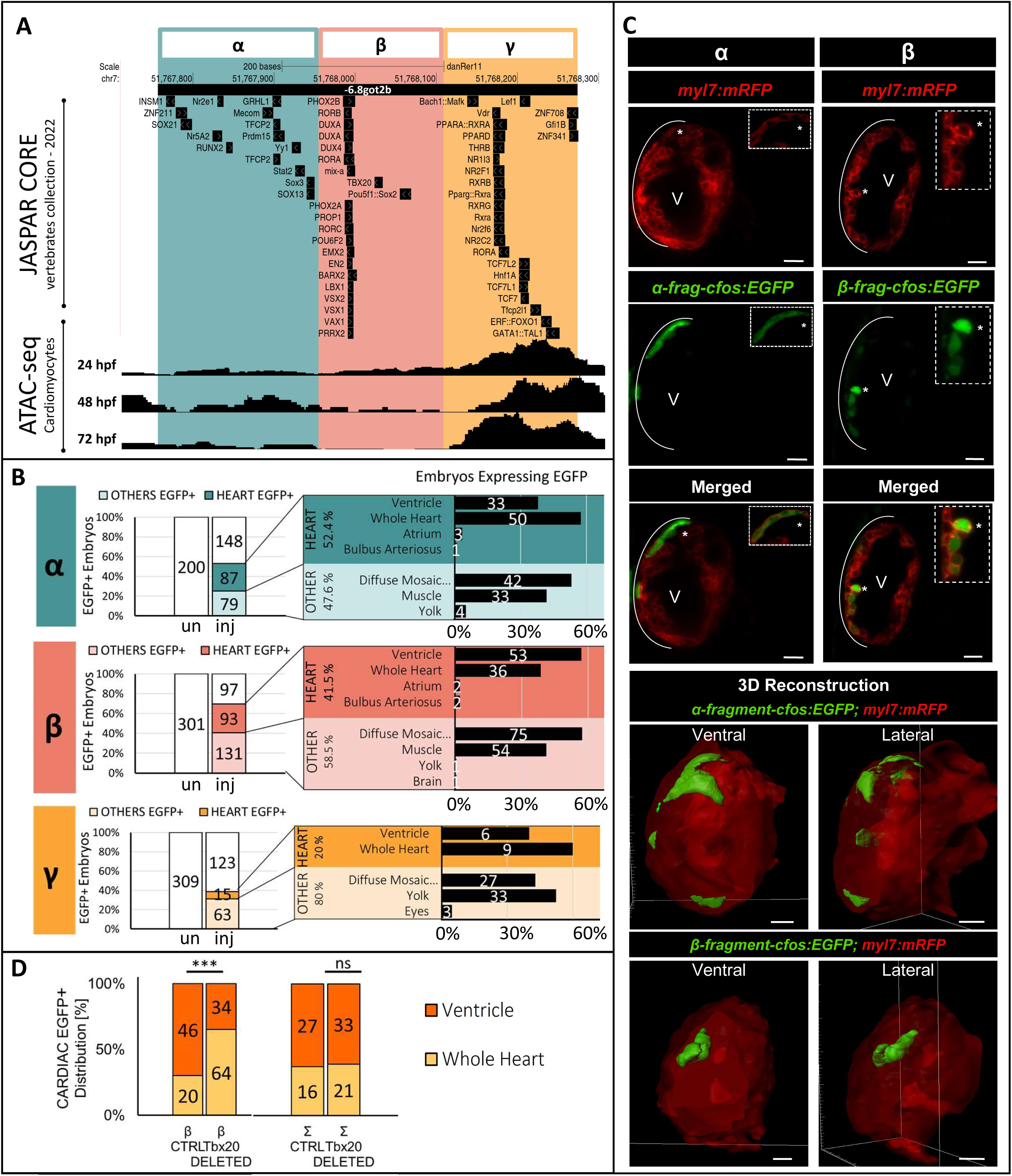
The core enhancer region of *-6.8got2b* confers specific expression in trabecular cardiomyocytes through Tbx20. **A**, transcription factor binding motifs (JASPAR CORE vertebrates collection 2022) and open chromatin region (ATAC-seq of whole cardiomyocytes^49^ within the *- 6.8got2b* region. **B**, quantification of reporter expression at various tissues driven by α, β, and γ enhancer fragments. Data acquired at 72 hpf from 3 independent experiments. **C**, visualization of EGFP expression driven by α and β fragments in *Tg(myl7:mRFP)* embryos and its 3D reconstruction. **D**, changes in reporter expression pattern upon deletion of Tbx20 motif from β and Σ regions. ***P < 0.001 (Fisher’s test).

**Table 1.** List of TF motifs found in −6.8got2b enhancer region and its respective subfragments tested.

| TFBMs | Related Signalling pathways form GeneCards®: The Human Gene Database |
| --- | --- |
| **α-fragment** |  |
| Nr2e1 | PIP3 activates AKT signalling. |
| RUNX2 | Bone development. |
| ZNF211 | Gene Transcription pathways. |
| TFCP2 | Signalling events mediated by HDAC Class I; Angiogenesis. |
| Yy1 | Notch signalling pathway, Sterol regulatory element-binding proteins signalling. |
| Stat2 | Cytokine Signalling in Immune system. |
| SOX13 | Signalling by WNT and ERK Signalling. |
| Sox3 | Signalling by WNT and ERK Signalling. |
| Prdm15 | FOXA1 transcription factor network; Nuclear estrogen receptor alpha network. |
| Nr5A2 | ESR-mediated signalling; Lipid metabolism. |
| GRHL1 | PPARA activates gene expression; Metabolism; Epithelial development. |
| INSM1 | Regulation of beta-cell development and Nervous system development. |
| Mecom | PIP3 activates AKT signalling; Chromatin organization. |
| SOX21 | ERK Signalling; Mesodermal commitment pathway. |
| **β-fragment** |  |
| PHOX2A | Sudden infant death syndrome (SIDS) susceptibility pathways. |
| PROP1 | Pituitary hormone regulation. |
| POU6F2 | Wilms Tumor 5 and Wilms Tumor And Radial Bilateral Aplasia associated Diseases. |
| BARX2 | Pre-implantation embryo. |
| PHOX2B | Sudden infant death syndrome (SIDS) susceptibility pathways and Neural crest differentiation. |
| mix-a | Not found. |
| RORC | Gene Transcription; Cytokine production by Th17 cells in Cystic Fibrosis. |
| RORB | Sweet Taste Signaling; Clock-controlled autophagy in bone metabolism. |
| VSX2 | Markers of kidney cell lineage. |
| PRRX2 | mammalian dermal regeneration; foetal skin development . |
| LBX1 | Key regulator of muscle precursor cell migration and is required for the acquisition of dorsal identities of forelimb muscles in mice. |
| VSX1 | expression of the cone opsin genes early in development. |
| EMX2 | Kidney development; Nervous system development. |
| DUX4 | RNA Polymerase I Promoter Opening; Nervous system development. |
| EN2 | Dopaminergic neurogenesis. |
| TBX20 | Cardiogenesis; Nervous system development. |
| VAX1 | Dorsoventral specification of the forebrain. |
| Pou5f1::Sox2 | Cardiac progenitor differentiation. |
| DUXA | Maternal to zygotic transition , Zygotic genome activation. |
| RORA | Circadian rhythm and lipid and glucose metabolism. |
| **γ-fragment** |  |
| PPARA::RXRA | PPAR signalling pathway; Lipid metabolism. |
| Pparg::Rxra | PPAR signalling pathway; Lipid metabolism. |
| THRB | Thyroid hormone signalling pathway. |
| TCF7L2 | Wnt signalling pathway. |
| PPARD | PPAR signalling pathway; Energy metabolism; Fatty acid metabolism, Pyruvate metabolism. |
| NR1I3 | Xenobiotic and endobiotic metabolism; Retinoic acid response. |
| TCF7 | Wnt signalling pathway. |
| Hnf1A | ERK Signalling; Regulation of beta-cell development. |
| Lef1 | Wnt signalling pathway. |
| TCF7L1 | Wnt signalling pathway. |
| NR2C2 | Repressor of nuclear receptor pathways including retinoic acid, retinoid X, vitamin D3, thyroid hormone, and estrogen receptors. |
| Tfcp2l1 | Cell morphogenesis; regulation of transcription by RNA polymerase II. |
| RXRG | PPAR signalling pathway; Lipid metabolism. |
| Nr2f6 | Hormonal responses. |
| RXRB | PPAR signalling pathway. |
| Rxra | PPAR signalling pathway. |
| ERF::FOXO1 | Extraembryonic ectoderm differentiation; ectoplacental cone cavity closure; chorioallantoic attachment; IL-9 Signaling Pathways and FOXO-mediated transcription. |
| GATA1::TAL1 | Erythropoietin signalling pathways. |
| Gfi1B | NF-kappaB Signalling , erythroid development and differentiation. |
| ZNF341 | JAK/STAT signalling pathway (cytokine signalling , development, immunity, and tumorigenesis). |
| Vdr | Vitamin D Receptor Pathway. |
| NR2F1 | Steroid/thyroid hormone receptor pathway for brain development. |
| ZNF708 | Breast Cancer Progression. |
| Bach1::Mafk | Nuclear events mediated by NFE2L2 and Cellular responses to stimuli. |
| RORA | Circadian rhythm and lipid and glucose metabolism. |

In order to further pinpoint the upstream regulators acting via the core enhancer region, we examined TF motif occurrence in this region. Fragment β harbors the binding motifs of 20 TFs involved in a wide range of developmental pathways and signalling cascades (**Table 1**). Among them is Tbx20, a TF previously implicated in mammalian cardiac ventricular wall trabeculation and maturation^56^. To explore its role in regulating reporter activity in trabecular cardiomyocytes, we deleted this motif from the vector Tol2:β*-fragment-cfos:EGFP* (**Supplemental Figure S6A**). In the controls injected with the vector carrying the unmutated β fragment, EGFP expression was predominantly ventricular (∼70% ventricle, ∼30% whole heart), whereas Tbx20 motif removal reversed the distribution (∼35% ventricle, ∼65% whole heart), indicating a loss of ventricular specificity (**Figure 4D, Supplemental Figure S6B**). To investigate potential compensatory effects, we deleted the Tbx20 motif from the full – *6.8got2b* enhancer sequence (here referred to as Σ). The expression pattern (∼63% ventricle, ∼37% whole heart) remained largely unchanged upon deletion of the Tbx20 motif, suggesting alternative mechanisms maintaining ventricular EGFP expression. (**Figure 4D, Supplemental Figure S6B**).

Taken together, all 3 fragments representing distinct regions of the *-6.8got2b* enhancer could drive cardiac expression to varying degrees, with the β fragment showing the strongest and most specific expression in trabecular cardiomyocytes in the ventricle. This pattern closely resembled that of the full enhancer region, indicating that the core regulatory activity of the *- 6.8got2b* enhancer resides within the genomic region represented by the β fragment. Furthermore, the requirement for the Tbx20 binding motif for maintaining trabecular-cardiomyocyte-restricted expression establishes Tbx20 as a key upstream regulator acting through the *–6.8got2b* enhancer to control ventricular trabeculae development.

### Evolutionary conservation of the *–6.8got2b* enhancer in humans

To establish the evolutionary conservation of the *-6.8got2b* enhancer, we applied direct sequence comparisons to search for putative regulatory elements harbouring similar sequences in evolutionarily close and distant species. Within Cyprinids, an element of 181 bp was found with 75.7% identity in the goldfish genome (*Carassius auratus).* Similarly, an element of 232 bp and 158 bp was found with 75.4% and 75.3% identity in golden-line barbel (*Sinocyclocheilus grahami*) and common carp (*Cyprinus carpio*) respectively (**Figure 5A**). These sequences were also found in synteny with the corresponding *got2b* gene in each species, which is one hallmark of enhancer-target gene regulatory conservation^57^. Beyond the Cyprinid family, direct sequence alignment with the spotted gar, an ancient fish which bridges teleosts to tetrapods^58^, did not identify any significant match. We then extended the search across multiple taxa comprising 14 orders with a total of 32 species, including the painted turtle which is a species with slowly evolving genomic sequence. This also yielded no significant matches. Pairwise whole genome alignment between zebrafish and human (https://genome.ucsc.edu/cgi-bin/hgTables?db=rheMac10&hgta_group=compGeno&hgta_track=chainNetHg38&hgta_ta ble=chainHg38&hgta_doSchema=describe+table+schema) revealing a 178 bp sequence at chr12:16,512,987-16,513,164 in the human genome (GRCh38/hg38) with the highest-scoring value for syntenic elements. This human element, from hereon designated as −0.17GOT2P4, covers 34.4% of the zebrafish *-6.8got2b*, with 48.7% identity (**Figure 5B**) and overlapped H3K27ac epigenetic mark for active enhancer in the ENCODE database (**Supplemental Figure S7**). The human element shares 17 TF binding motifs with *–6.8got2b,* among which, 47% (8 out of 17) overlap with motifs found in the γ fragment of the zebrafish enhancer, 47% (8 out of 17) overlap with motifs in the β fragment, and 6% (1 out of 17) overlap with a motif in the α fragment (**Figure 5C**). These include TCF7, TCF7L2, LEF1, TCF7L1, and RORA, present within the β and γ fragments. However, unlike −*6.8got2b* and *got2b* relationship in the zebrafish, the human *-0.17GOT2P4* did not retain syntenic conservation with *GOT2* which was located in chromosome 16. Intriguingly, *GOT2 pseudogene 4 (GOT2P4)*, a pseudogene which shares 90% of the complete protein coding sequence of *GOT2*, was found immediately downstream (0.17 kb) of the human element. *GOT2P4* lacked all 9 introns found in its functional counterpart at chr16, suggesting that it arrived in the current genomic loci through retrotransposition. The syntenic relationship between the human element and *GOT2P4* pseudogene is conserved among primates, suggesting a relatively recent transposition event (**Supplemental Figure S7B**).

**Figure 5.**
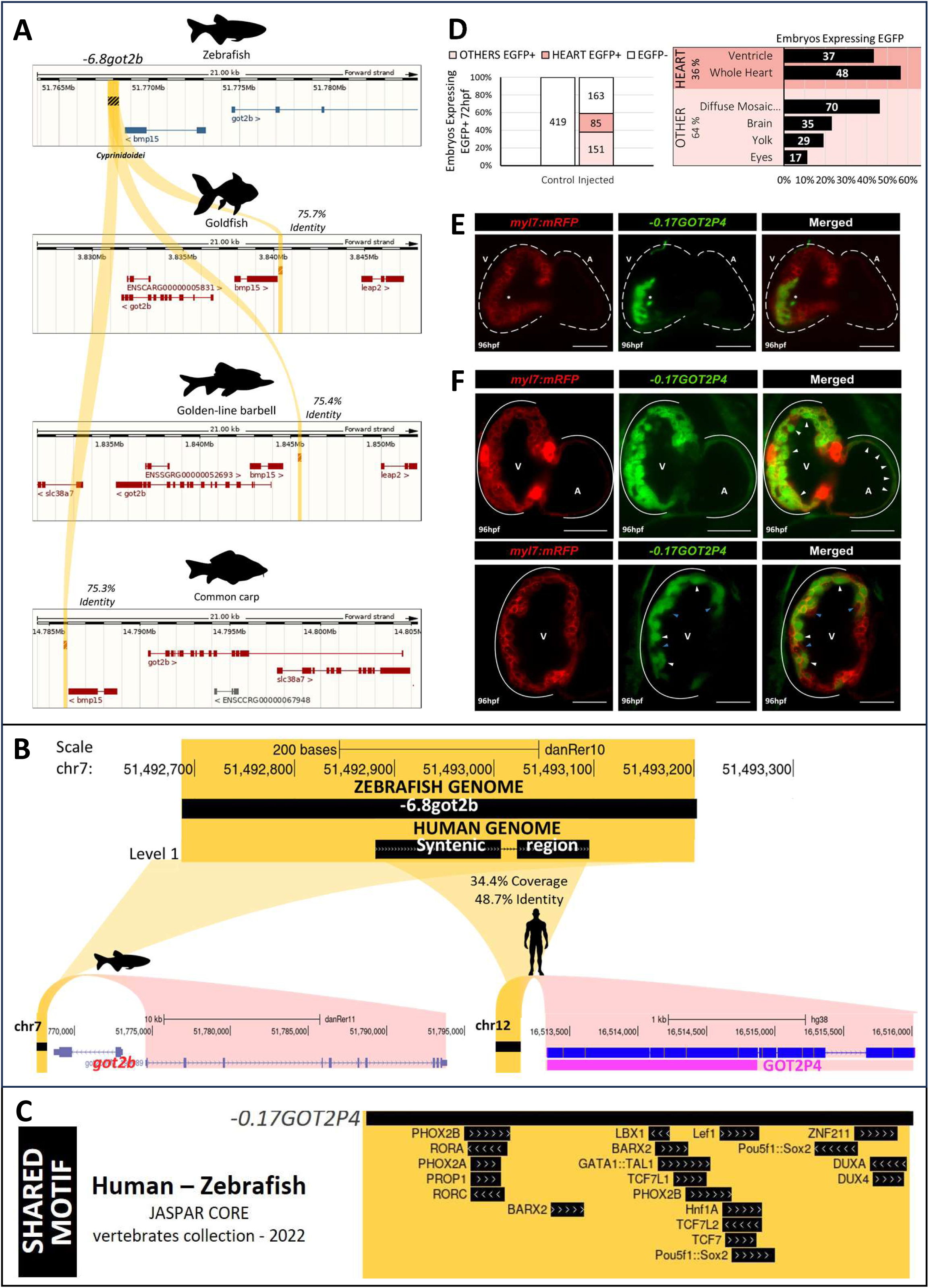
Conservation analysis of *-6.8got2b* enhancer. **A**, visualization of the *–6.8got2b* enhancer in the zebrafish genome, with corresponding conserved sequences in the goldfish (*Carassius auratus*), golden-line barbel (*Sinocyclocheilus grahami*), and common carp (*Cyprinus carpio*) highlighted in yellow. **B**, zebrafish - human sequence alignment revealed –0.17GOT2P4 in proximity to the GOT2 pseudogene 4 in human. **C**, the –0.17GOT2P4 human region with shared Human-Zebrafish TF binding motifs from JASPAR CORE vertebrates collection. **D**, quantification of reporter expression at various tissues driven by the human –0.17GOT2P4. Data acquired at 72 hpf from 3 independent experiments. **E**, light sheet fluorescent image of the heart in *Tg(myl7:mRFP)* embryos injected with the *– 0.17GOT2P4 cfos-EGFP-Tol2* construct at 96 hpf. Scale bar = 50 μm; A = Atrium; V = Ventricle. **F**, light sheet fluorescent image of the heart in the F1 individuals carrying the *Has.-0.17GOT2P4-cfos:EGFP* construct showing two distinct patterns of EGFP expression in the whole heart (top) and subsets of ventricular cardiomyocytes (bottom). White arrows indicate EGFP+ cardiomyocytes, while the blue arrow marks cardiomyocytes lacking EGFP expression. Scale bar = 50 μm; A = Atrium; V = Ventricle.

To establish functional conservation between *–6.8got2b* and *–0.17GOT2P4*, we tested the ability of the latter to drive reporter expression in the enhancer activity assay. A region of 284 bp (chr12:16,665,875-16,666,158; GRCh37/hg19) was cloned into the vector and the resulting *-0.17GOT2P4-cfos-EGFP-Tol2* construct was microinjected into zebrafish embryos at the one-cell stage. Out of 399 injected embryos, 59% (236 out of 399) exhibited EGFP reporter expression at 72 hpf, among which, 36% (85 out of 236) showed expression primarily in the heart (**Figure 5D**). Within this group, the most common pattern was a uniform expression throughout the entire heart, accounting for 56.5% of all embryos expressing EGFP in this organ (48 out of 85). The second most common expression pattern was observed in the ventricle, present in 43.5% of embryos (37 out of 85). The EGFP-expressing cells in the ventricle overlapped with the cardiomyocyte RFP signal from Tg(*myl7*:mRFP) and are localized at the outer curvature of the ventricle, within the compact wall as well as the deeper trabecular cells (**Figure 5E**). A stable transgenic line, *Tg(Has.-0.17GOT2P4-cfos:EGFP)*, was subsequently generated by raising the 37 embryos exhibiting EGFP+ heart expression, of which 32 survived to adulthood. Nine individuals were subsequently outcrossed with wild-type and their offsprings (F1) were observed at 96 hpf. All of them exhibited germ line transmission and consistently expressed the EGFP reporter in the heart. The offspring from two F0 fish were characterized by EGFP expression in the whole heart myocardium, while the offspring from other 2 F0 had EGFP signal in a subset of ventricular cardiomyocytes (**Figure 5F**).

### Transcriptome profiling identifies molecular signatures distinguishing trabecular from compact cardiomyocytes

The specificity and early onset of reporter expression pattern driven by the *-6.8got2b* enhancer allows the identification and isolation of trabecular cardiomyocytes prior to the appearance of observable trabecular structure. We therefore leveraged this system to investigate molecular features distinguishing trabecular from compact cardiomyocytes making up the wall of the ventricular chamber. We outcrossed the *Tg(−6.8got2b:EGFP)* line with *Tg(myl7:mRFP)* to obtain embryos in which compact cardiomyocytes are marked by RPF and trabecular cardiomyocytes by both RFP and EGFP. FACS isolation of these populations at 48 hpf followed by transcriptome profiling revealed a distinct transcriptional signature of trabecular cardiomyocytes compared to that of the compact layer (**Figure 6A**). Differential expression analysis identified genes selectively enriched in trabecular cardiomyocytes, including *got2b*, highlighting candidate molecular markers defining early trabecular identity (**Figure 6B, Supplemental Table S1**).

**Figure 6.**
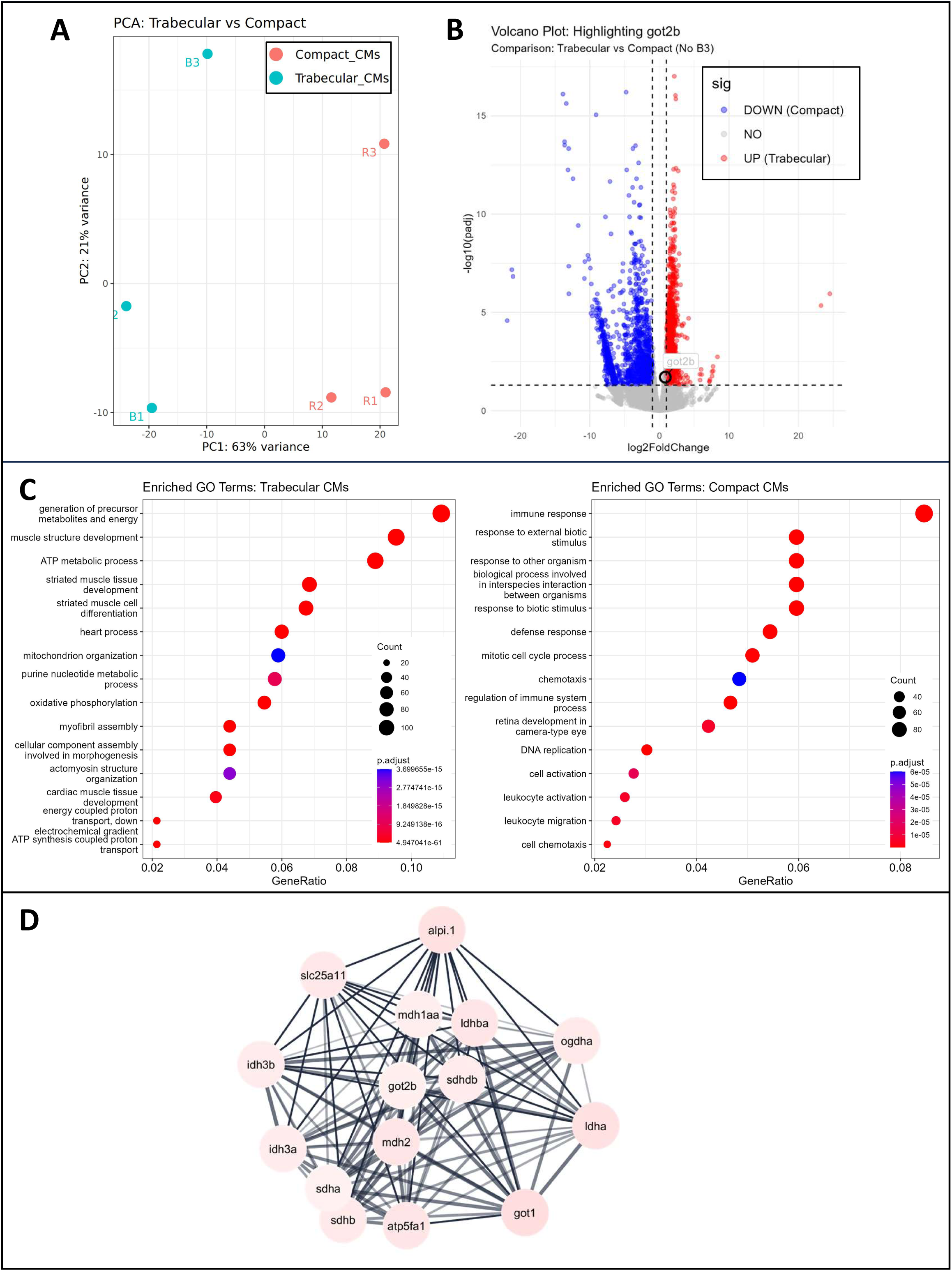
Transcriptome profile of trabecular and compact cardiomyocytes. **A**, principal component analysis (PCA) showing the separation of trabecular (B1–B3) and compact layer (R1–R3) cardiomyocytes based on global gene expression profiles. **B**, volcano plot highlighting differentially expressed genes between trabecular and compact cardiomyocytes. **C**, Gene Ontology (GO) terms significantly enriched in trabecular versus compact cardiomyocyte populations. **D**, protein–protein interaction network centered on Got2b, revealing connections to metabolic enzymes and mitochondrial pathways.

The trabeculae-enriched genes can be broadly grouped into contractile or sarcomeric components and developmental regulatory factors, reflecting both functional and morphogenetic specialization. Contractile genes include *tnnt2b*, which encodes a cardiac troponin T isoform essential for sarcomere function and heartbeat initiation, and *tnni4a*, encoding a troponin I subunit mediating calcium-dependent actin–myosin regulation. Together, these genes reflect the contractile identity of trabecular cardiomyocytes^59^. Trabecular cardiomyocytes also overexpress *csrp3* (Muscle LIM protein), encoding a Z-disc–associated factor critical for striated muscle integrity and limiting trabecular growth in zebrafish^60^, as well as *lrrc10*, encoding a cardiac-specific regulator of embryonic myocardial contraction^61^, and *pln1*, encoding a sarcoplasmic reticulum protein modulating calcium reuptake via SERCA inhibition and thereby fine-tunes cardiomyocyte contractility^62^. Several genes implicated in cardiac development and signaling are also enriched in trabecular cardiomyocytes. These include *tbx20,* which encodes the upstream regulator of *–6.8got2b* enhancer, and *tbx5a*, encoding a T-box transcription factor essential for chamber identity and morphogenesis. Components of growth factor signaling are also enriched, including *bmp10l* and *bmp7* which regulate trabecular myocardial growth and cardiomyocyte proliferation^63,64^ as well as *erbb4*, the receptor for Neuregulin 1, which, together with *erbb2*, mediates endocardial signalling essential for the onset of trabeculation and the specification of trabecular cardiomyocytes^39,65^, while *wnt6b* prevents excessive myocardial expansion^66^, consistent with its role in chamber formation. In addition, *nppa*, a well-established marker of embryonic trabecular myocardium, actively contributes to chamber morphogenesis, as dysregulation leads to abnormal chamber expansion and cardiac malformations^67^.

Gene Ontology (GO) enrichment analysis shows that terms related to cellular metabolism and energy production (such as “generation of precursor metabolites and energy”, “ATP metabolic process”, and “oxidative phosphorylation”), as well as “muscle structure development”, were significantly overrepresented among genes enriched in trabecular cardiomyocytes (**Figure 6C**, **Supplemental Table S2**). On the other hand, genes enriched in compact cardiomyocytes were associated with less significant enrichment for GO terms associated with basic cellular functions (**Figure 6C**, **Supplemental Table S2**). Reactome pathway enrichment analysis of trabecular-enriched genes highlighted pathways involved in mitochondrial processes and glucose metabolism (**Supplemental Figure S8**, **Supplemental Table S3**), further supporting a metabolic specialization of trabecular cardiomyocytes at early developmental stages. In addition, protein–protein interaction network analysis centered on *got2b* connects it to metabolic enzymes and mitochondrial pathways, placing it within the broader metabolic network active in trabecular cardiomyocytes (**Figure 6D**, **Supplemental Figure S9**). Collectively, these data align with the expected high metabolic demand of the developing trabeculae, supporting that trabecular cardiomyocytes are already transcriptionally primed for elevated energy production prior to overt trabecular morphogenesis.

## Discussion

Trabecular cardiomyocytes are essential for ventricular function, yet the regulatory programs driving their development remain poorly defined. Here, we identify the *-6.8got2b* enhancer as an early and specific regulator of trabecular fate that primes cardiomyocytes for elevated metabolic activity before structural trabeculation becomes apparent. Through its target *got2b*, this conserved –*6.8got2b–got2b* axis operates downstream of Tbx20 to couple transcriptional identity with metabolic state, revealing an unrecognized layer of transcriptional regulatory control orchestrating early trabecular specification and ventricular patterning. While recent advances in imaging and computational reconstruction have provided detailed three-dimensional views of trabecular architecture^68,69^, the molecular distinction between compact and trabecular cardiomyocytes, particularly at early developmental stages, has remained elusive. Notably, the *–6.8got2b* -driven expression is detectable from 48 hpf, establishing it as the earliest known specific marker distinguishing trabecular from compact cardiomyocytes and enabling the resolution of fate specification before morphogenesis.

Linking enhancers to their direct target genes remains a central challenge in regulatory biology, as some act over long genomic distances^17,70^ and may function combinatorially, complicating target identification^71,72^. Nonetheless, studies show that most enhancers preferentially regulate genes within their local chromosomal neighbourhood, often constrained by topologically associating domains (TADs)^10–12,73^. Notably, *–6.8got2b* and *got2b* reside within the same predicted TAD, supporting a structurally permissive chromatin environment for regulatory interaction. Multiple independent lines of evidence support *got2b* as the primary target gene of the *–6.8got2b* enhancer. Among neighbouring genes within the TAD, *got2b* exhibits consistently high expression in cardiomyocytes that increases through development^49^. Importantly, our single-cell transcriptomic analysis of the entire zebrafish heart^54^ confirms robust and selective expression of *got2b* within ventricular cardiomyocytes, closely mirroring the spatial and temporal activity of the enhancer. In contrast, other genes within the same genomic interval show little or no ventricular expression. Together, genomic proximity, shared TAD architecture, and concordant single-cell expression patterns strongly support a direct regulatory relationship between –*6.8got2b* and *got2b*.

Zebrafish *got2b* is an orthologue of the human GOT2 gene^74^, encoding the mitochondrial glutamic-oxaloacetic transaminase 2, a core component of the malate–aspartate shuttle that supports ATP production (reviewed in Safer 1971^75^; Scholz et al. 1998^76^). Within the broader context of cardiac trabeculation, Got2b likely supports the high energy demands associated with cardiomyocyte delamination and morphogenesis. Consistent with this, comparison of trabecular and compact cardiomyocytes transcriptomes reveals a distinct trabecular signature characterized by enhanced metabolic and cell adhesion functions, reflecting elevated energy requirements and dynamic cellular behaviour. These results align with studies showing that ErbB2-dependent trabeculation in zebrafish requires glycolytic activation^36^, and that cardiomyocytes undergo metabolic changes during regeneration, shifting from oxidative metabolism to a more glycolytic state^77^, highlighting the central role of metabolic adaptation in cardiomyocytes. Our findings extend these observations by demonstrating that metabolic specialization is established transcriptionally before overt trabecular morphology emerges, indicating that trabecular cardiomyocytes are pre-patterned for elevated energy production prior to structural differentiation.

Although *got2b* is expressed in ventricular, muscle, and neural tissues^78^, the *–6.8got2b* enhancer drives expression specifically in trabecular cardiomyocytes and a restricted domain of the choroid plexus. This restricted activity suggests that *–6.8got2b* regulates a subset of *got2b* expression, exemplifying how enhancers fine-tune gene expression and partition pleiotropic gene functions into discrete spatiotemporal domains^73,79^. This property makes enhancers powerful tools for resolving and marking discrete cellular identities. Dissection of the enhancer revealed three subregions within the *–6.8got2b* enhancer that exhibit distinct activities in driving EGFP expression across myocardial subpopulations, consistent with modular enhancer architecture and combinatorial transcription factor logic^80,81^. Notably, although the β fragment most closely recapitulated the activity of the full-length enhancer, open chromatin profiling of bulk cardiomyocyte^49^ indicated accessible region within the γ fragment. This discrepancy likely reflects the underrepresentation of trabecular cells at early stages of heart development. Ongoing single-cell chromatin analyses will further resolve enhancer accessibility differences between compact and trabecular cardiomyocytes.

The *–6.8got2b* enhancer is highly conserved among Cyprinids with approximately 75% sequence identity, consistent with selective constraint on key developmental regulatory elements^19^. Conserved TF motifs found include those associated with energy, fatty acid, protein, and pyruvate metabolism, as well as Wnt and Hippo signalling pathways, reinforcing its role in coordinating metabolic state with trabecular morphogenesis. In contrast, the absence of detectable sequence conservation with distant species such as spotted gar indicates that *– 6.8got2b* may represent a lineage-specific regulatory adaptation. This underscores the evolutionary plasticity of enhancers, which can remain highly constrained within clades while diversifying to generate lineage-specific regulatory architectures^82,83^.

A human genomic element, –0.17GOT2P4, sharing partial sequence similarity with *– 6.8got2b*, drove similar cardiac expression in zebrafish, indicating conservation of regulatory logic across vertebrates. This human element retains key transcription factor motifs clustered within the core functional region of the zebrafish enhancer, supporting preservation of motif architecture despite sequence divergence. Intriguingly, –0.17GOT2P4 resides near the pseudogene GOT2P4 instead of the functional GOT2 locus. GOT2P4 lacked all 9 introns of its functional counterpart, suggesting that it arrived in the current genomic loci through retrotransposition from the ancestral coding gene^84^. The absence of a second GOT2 copy in non-teleost vertebrates suggests that the two zebrafish paralogs arose by teleost-specific whole genome duplication, supporting an argument that the human pseudogene represents a retrotransposed remnant rather than a degenerated ancestral paralog. The proximity of – 0.17GOT2P4 to this pseudogene raises the possibility that enhancer logic can persist independently of its original coding locus, and that genomic rearrangements may reposition regulatory elements into new contexts while maintaining functional capacity. Whether the conserved enhancer architecture was independently retained near the pseudogene or co-mobilized during retrotransposition remains unresolved. Together, our findings highlight the complexity and evolutionary plasticity of enhancer-gene relationships and illustrate how regulatory elements can be retained, repurposed, or uncoupled from their ancestral targets during vertebrate genome evolution^82,83^.

## Supplementary Material

### Supplemental Figures

**Supplemental Figure S1.** Schematics of the zebrafish *–6.8got2b* region showing sequence variations in the region isolated from AB and TL wild-type strains: a 2-bp gap at position chr7:51767950-51767951 (GRCz11/danRer11), and a A to G variation at position chr7:51768224-51768224 (GRCz11/danRer11).

**Supplemental Figure S2.** Whole-embryo reporter expression pattern of the Tg(*-6.8got2b-cfos:EGFP)* at 96 hpf encompassing cardiac ventricle and a restricted domain at the choroid plexus.

**Supplemental Figure S3.** Transgenic reporter expression in Tg(*-6.8got2b-cfos:EGFP)* x Tg*(kdrl:mCherry)* outcross, showing distinct *-6.8got2b*-driven EGFP expression domain and the endocardial mCherry.

**Supplemental Figure S4.** The ratio of spatial spread of trabecular and myocardial cells for selected angular sectors for indicated conditions, CTRL (black, n=3, 32 Z-slices), SCRMB (gray, n=3, 50 Z-slices), INJ (red, n=5, 74 Z-slices), mean ± 95% CI. ns (not significant); one-way ANOVA with the Bonferroni’s post hoc test.

**Supplemental Figure S5. A**, Hi-C analysis of muscle cells showing topologically associating domain ^53^ containing several gene loci within approximately 100 kb region up- and downstream of *–6.8got2b*. **B**, single-cell RNA-seq data of the whole heart ^54^, showing *got2b* expression in the ventricular cardiomyocyte cluster. **C-E**, no significant expression of *hdac8* and *slc38a7* and *leap2* was observed in the single-cell RNA-seq data.

**Supplemental Figure S6. A**, schematics of the –6.8got2b enhancer β fragment (blue) and full-length (Σ, red), showing deletion of the Tbx20 motif and its confirmation by Sanger sequencing. **B**, quantification of reporter expression in various structures driven by the wild-type enhancer regions and those carrying the Tbx20 motif deletion.

**Supplemental Figure S7. A**, epigenetic profiling shows H3K27ac and open chromatin regions in the human –0.17GOT2P4. **B**, visualization of the –0.17GOT2P4 region and GOT2P4 pseudogene in the human genome, with corresponding conserved sequences in *Gorilla gorilla*, *Pan troglodytes*, and orangutan *Pongo pygmaeus* highlighted in yellow and brown.

**Supplemental Figure S8.** Reactome pathway enrichment analysis of trabecular-enriched genes.

**Supplemental Figure S9.** Protein–protein interaction network analysis showing the broader metabolic network enriched in trabecular cardiomyocytes.

**Supplemental Table S1.** List of differentially expressed genes in trabecular vs compact cardiomyoctes.

**Supplemental Table S2.** Enrichment of gene ontology terms.

**Supplemental Table S3.** Reactome Pathway analysis.

**Supplemental Table S4.** List of all genes expressed in RNA-seq with their expression values.

## Methods

### Zebrafish maintenance

Wild-type (AB and TL strain) and transgenic line, Tg(*myl7:EGFP-Hsa.HRAS*)^s883^ ^1^, Tg(*myl7:mRFP*) ^2^, Tg(*kdrl:Hsa.HRAS-mCherry*)^3^, Tg(*–6.8got2b-cfos:EGFP*) and Tg(*Has.–0.17GOT2P4-cfos:EGFP*) were maintained in the zebrafish core facility of the International Institute of Molecular and Cell Biology (license no. PL14656251). Embryos were staged according to standard morphological criteria^4^ and maintained at 28°C in egg water (60 mg/liter Instant Ocean Salts) with a 14 hr light/10 hr dark cycle. All zebrafish husbandry was performed under standard conditions in accordance with institutional national ethical and animal welfare guidelines. 1-phenyl 2-thiourea (PTU) was added at 24 hpf to prevent pigmentation. Embryonic stages are given as hours post-fertilization at 28 °C.

### *in vivo* enhancer assays and screening

Zebrafish transgenic assays and screening were performed following previously established procedures^5,6^. Briefly, the genomic region identified as putative enhancer was amplified by PCR and cloned into *Kpn*I and *Bam*HI sites of the Tol2-EGFP reporter vector containing a minimal promoter from the mouse *cFos* gene. The vector construct containing the putative enhancer was then sequenced to verify the correct insertion. Transposase mRNA was synthesised by in vitro transcription using mMESSAGE mMACHINE® Kit (Life Technologies) and purified using RNA Clean & Concentrator-5 (Zymo Research). A total of 20 pg of the circular reporter vector and 50 pg of transposase mRNA were co-injected into 1 cell stage embryos and assayed for EGFP expression from 24 hpf.

### Light-sheet fluorescence microscopy and 3D reconstruction

Embryos were anaesthetised using Tricaine (MS-222) at desired developmental stages, embedded in a 1 mm inner-diameter glass capillary containing 1.5% low-melting agarose (LMA) in Tricaine (MS-222). The microscope chamber containing the sample holder and the capillary was filled with 15X Tricaine. Image acquisition was performed using a Zeiss Z.1 light-sheet microscope with a W Plan-Apochromat 40×/1.0 objective. Z-stacks (3.83 μm thickness, 60 ms exposure time) were saved in LSM format and then processed using ZEN software (Zeiss). For each Z-stack, maximum-intensity projections were generated. 3D images were processed using Imaris software.

### 3-Dimensional Reconstruction

Each Z-stack data set from the light-sheet was converted using the Imaris File Converter and uploaded to Imaris software 8.3 version. Representative images from control conditions were used to set up the batch protocol. It includes creating a surface object for each channel using “Surfaces” module. The settings mask for EGFP and mRFP signals are listed in **Table 2**. The parameters were saved and applied to each condition.

**Table 2.** Parameters of Surface object in Imaris software.

| Parameter | EGFP | mRFP |
| --- | --- | --- |
| Smooth surface | True | True |
| Surface Grain Size | 0.5 $\mu\text{m}$ | 1 $\mu\text{m}$ |
| Diameter of largest sphere | 0.7 $\mu\text{m}$ | 0.7 $\mu\text{m}$ |
| Classify surfaces | Distance to image border XYZ | Distance to image border XYZ |

Morphological parameters for each surface object were then measured, including: “Area”, expressed in square micrometres (µm²); “Volume”, expressed in cubic micrometres (µm³); and “Sphericity”, a measure of how spherical an object is, ranging from 0 to 1. A value of 0 suggests that the object is highly irregular or elongated, while a value of 1 indicates a perfect sphere. These parameters were exported as CSV files for calculations and statistical analysis.

### 2-Dimensional spatial analysis

To complement the 3-Dimensional reconstruction we developed workflow for direct quantification of myocardium structure using individual Z-stack images (slices). For each Z-stack data set for 120 hpf we selected every fourth slice along z-axis, this gave 74 slices in total for −6.8got2b loss of function (n = 5 Injected) and 82 slices for controls (n = 6, 3 Control, 3 Scrambled). The slices were imported separately for GFP and RFP signals. The data was analysed with a self-written custom script in Wolfram Mathematica v14.3. The background was removed with the default RemoveBackground function. The data was manually trimmed to match minimal rectangular region containing the heart and results were binarized using default settings to positions of GFP/RFP pixels. Next, the position of the center of the heart was identified as either the center of mass of all pixels for a given Z-stack slices for joined GFP and RFP signals, or through manual adjustment (**Figure 3F**). The center of the heart was used as origin of the polar coordinates, mapping the (x,y) pixel positions to their respective angular and radial coordinate, (ϕ, r). The data was then binned into 36 angular sectors spaced every Δϕ = π/18 radians. To avoid regions with empty sectors, for each Z-stack, the sectors of interest were selected as a range of ϕ (**Figure 3F**). Further, only sectors with at least 10 pixels in each channel were used to compare the positions of GFP and RFP pixels. The wall thickness was estimated as a difference between maximal and minimal radius of any pixel (GFP and RFP signals pooled together) within each sector (**Figure 3G**). The average radial distance of trabecular and compact myocardial cells from the center of the heart was calculated as the mean of the radial coordinate of pixels of respective RFP and GFP pixels within each sector, thus positive shift corresponds to EGFP+ cells shifted towards the center of the heart relative to RFP signal (**Figure 3H**). The spatial spread of trabecular and myocardial cells was estimated as the standard deviation of respective GFP and RFP pixels within each sector (**Supplemental Figure S4**). The means reported in the main text were obtained by pooling together estimates for each sector and each slice and averaging per condition.

#### Chemical treatments

Larvae were exposed to Tyrphostin (AG1478, Sigma-Aldrich) at a concentration of 2 μM, following the procedure described by Liu et al., 2010^7^. Egg water containing 0.4% DMSO was utilized for chemical solubilization, and control larvae were incubated in 0.4% DMSO. The treatment was administered from 68 to 96 hpf.

#### gRNA design and Cas9 ribonucleoprotein complex assembly

gRNA sequences were designed based on the principles described in a previously established protocol^8^ using the CHOPCHOP algorithm with default settings for CRISPR/Cas9 knock-out method with efficiency score option^9^. Two gRNAs were designed to target upstream and downstream of the enhancer sequence and three different exons of the *got2b* gene. A scramble gRNA which does not have a complementary sequence in the zebrafish genome to direct a Cas9 nuclease to cut the genome was used as a negative control (**Table 3**). Synthetic gRNAs (sgRNAs) were ordered from Synthego at 1.5 nmol scales and dissolved in 15 μl of water to reach 100 μM stock concentration. A mix containing two or three sgRNAs in equal amounts was prepared to reach 57 μM (28.5 μM each). In parallel, a solution of a scramble sgRNA at 57 μM was prepared. Cas9 protein,(Alt-R™ S.p. Cas9 Nuclease V3, cat. no. 1081059, IDT), was diluted to 57 μM using Cas9 buffer (20 mM Tris-HCl, 600 mM KCl, 20% glycerol) in accordance with a previously documented protocol.

**Table 3.**
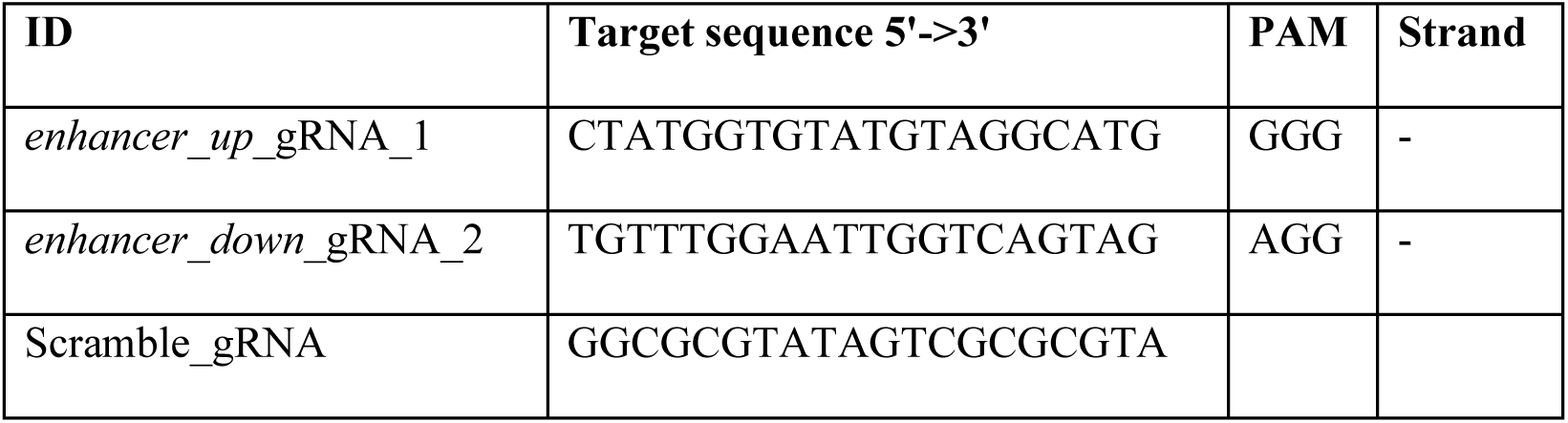
List of RNA guides for CRISPR-Cas9.

To test the nuclease activity of the sgRNA-Cas9 complex, *in vitro* assays were carried out on polymerase chain reaction (PCR) products spanning the CRISPR-targeted regions amplified by the following primer pair: Forward – ACTGCTGGATATACTGTTGTCC and Reverse – TCCATGCACCCGAATTGATG. The cleavage reaction (10 µL) was prepared for each targeted region by mixing the following components: 1 µL RNP (28.5 μM) as previously described, 2 µL PCR product, 7 μL ddH_2_O. The mix was incubated for 37 °C for 2-6 h followed by proteinase treatment at 65 °C for 10 min to release the DNA from the Cas9 endonuclease. The reaction was analysed using gel electrophoresis with 2% agarose gel. *In-vitro* DNA cleavage by sgRNAs-Cas9 nuclease was effective, generating distinct fragments in relation to single or multiple guide efficacy.

#### CRISPR/Cas9 mutagenesis

One day prior to scheduled injections, respective sgRNA mixes targeting *-6.8got2b* or *got2b* loci was combined with previously diluted Cas9 protein to reach a final concentration of 28,5 μM ribonucleoprotein complex (RNP) concentration (molar ratio 2:1, 114 μM sgRNA mix : 57 μM Cas9). Scramble gRNA-Cas9 with a final concentration of 28,5 μM ribonucleoprotein complex (RNP) (molar ratio 1:1, 57 μM scramble sgRNA : 57 μM Cas9) was also prepared as control. The RNPs were incubated at 37 °C for 5 min then cooled on ice and stored in 4 °C until microinjection. One nanoliter of each RNP complexes was injected into the yolk of one-cell stage embryos. Uninjected siblings and those injected with a scrambled guide were used as control conditions. Observation was performed at 96 hpf when the trabeculae were fully developed and clearly visible. For F0 knockdown analysis, those showing a reduction in EGFP signal were genotyped to confirm the genetic modification. Quantitative morphological analysis was performed to measure the volume and shape of trabecular structures. To generate stable mutant lines, injected embryos were raised to adulthood and outcrossed with wild-type zebrafish to identify founders. To genotype, caudal fin clippings were collected from tricaine methanesulfonate (MS-222) anaesthetised adults. Genomic DNA extraction was performed ^10^ and amplified by PCR using primers flanking region.

### *-6.8got2b* enhancer alignment

Using the sequence *-6.8got2b* enhancer as a query, BLASTN searches were conducted at ensembl.org with a search sensitivity set to distant homologies^11^. Candidate BLAST hit regions were manually inspected for their location in relation to neighbouring genes. Local alignment tools Water (EMBOSS) were used to assess the percentage identity^12^. The UCSC genome browser’s pair-wise whole genomic alignment tracks (comprising chain and net alignments “Human Chain/Net - Human (Dec. 2013 (GRCh38/hg38)) were used to determine the syntenic conservation of the *-6.8got2b* enhancer in Human^13^. This dataset shows net tracks indicating the best human/zebrafish chains for every part of the zebrafish genome, highlighting syntenic regions and potential orthologs, as well as for studying genome rearrangement. The top-level (level 1) chains, which are the largest and highest-scoring chains that span this region, were considered. The genomic landscape within 3 kb of the - 0.17GOT2P4 region, using HUGO Gene Nomenclature track^14^ filtered by pseudogene type and Retroposed Genes V9 track, including pseudogenes (https://genome.ucsc.edu/cgi-bin/hgTrackUi?db=hg38&g=ucscRetroAli9). 19 Additional custom tracks were employed, including H3K27ac epigenetic marks (indicating active enhancers), and ATAC-seq data from cardiac tissue from various human samples. The H3K27ac signals included those from cardiac muscle cells (The ENCODE Project Consortium 2012; ENCFF446JBL_ENCFF852WXE_ENCFF190PRO_ENCFF190PRO H3K27), the left ventricle (The ENCODE Project Consortium 2012; ENCFF254JZR_ENCFF491UMQ_ENCFF556RUC_ENCFF491UMQ), and the atrium (The ENCODE Project Consortium 2012; ENCFF958IXF_ENCFF983HIO_ENCFF983HIO).

ATAC-seq data included samples from the left ventricle of a 51-year-old female (The ENCODE Project Consortium 2012; ENCSR117PYB/ENCFF148ZMS), the left ventricle of a 53-year-old female (The ENCODE Project Consortium 2012; ENCSR851EBF/ENCFF996TBQ), and the coronary artery of a 51-year-old female (The ENCODE Project Consortium 2012; ENCSR584AXZ/ENCFF352UTD). Track parameters: Track height 60 pixels from a range of 8-128, data view scaling group auto-scale, windowing function mean+whiskers.

#### Isolation of cardiomyocytes

Whole heart tissues from 48 hpf zebrafish embryos from *Tg(–6.8got2b-cfos:EGFP)* x *Tg(myl7:mRFP)* outcross were isolated following the protocol from^15^ with minor adjustments. Briefly, 700-1000 embryos were dechorionated with pronase (20mg/mL) digestion, washed with ice cold egg water, and dissociated in Gibco Leibovitz’s L-15 Medium (Invitrogen) supplemented with 10% fetal bovine serum (Sigma-Aldrich). Batches of 200 embryos were mechanically dissociated, filtered sequentially through 70 μm (VWR 732-2758) and 40 μm (VWR 732-2757) nylon cell strainers, and intact hearts were manually collected using a low-binding pipette tip under a fluorescent stereomicroscope. The collected hearts were transferred to a 2 mL low-binding Eppendorf tube containing 1 mL of medium and were dissociated according to the protocol described by Bresciani et al., 2018^16^. Heart tissues were enzymatically dissociated in 0.25% Trypsin-EDTA (Gibco) in PBS supplemented with 80 μL of collagenase (100 mg/mL) (Sigma-Aldrich) at 30°C for 30 minutes and quenched with ice-cold medium. Cells were washed, filtered through a 35 μm cell strainer, and viability assessed by trypan blue exclusion on countess automated cell counter (ThermoFisher, USA). LIVE/DEAD Violet Fixable Stain (ThermoFisher, USA) was added prior to fluorescence-activated cell sorting on the CytoFlex SRT (Beckman Coulter, USA). mRFP⁺EGFP⁻ (compact) and mRFP⁺EGFP⁺ (trabecular) cardiomyocytes were isolated directly into TRIzol™ reagent (Invitrogen). Altogether, samples from three independent biological replicates from different experiments were collected.

#### RNA-seq and bioinformatics analysis

RNA was isolated using the Direct-zol™ RNA Miniprep Kit (Zymo Research) according to the manufacturer’s instructions. RNA concentration were measured using the Quantus™ Fluorometer (Promega, USA), and RNA integrity were assessed on the RNA 6000 Pico chip with the Agilent Bioanalyzer 2100 system (Agilent, USA). cDNA synthesis and RNA-seq library preparation were performed using the NEBNext® Single Cell/Low Input RNA Library Prep Kit for Illumina (New England Biolabs, E6420). Libraries were sequenced on the Illumina NovaSeq 6000 platform at a depth of 20 million paired-end (2 x 100bp) reads per sample. Raw paired-end sequencing reads were assessed using FastQC tool. Subsequently, fastp (v0.20.1) was employed to trim adapter sequences and remove low-quality bases. The pre-processed reads were aligned to the *Danio rerio* reference genome (Build GRCz11/danRer11) using STAR (v2.7.10a). Duplication metrics were calculated using Picard MarkDuplicates (v2.27.4), however, PCR duplicates were retained in the final alignment to keep the quantitative dynamic range of the library. One trabecular cardiomyocyte sample (B3) was generated from extremely low RNA input. Although cDNA amplification was possible, subsequent RNA-seq analysis revealed that this sample behaved as an outlier relative to the other replicates, consistent with amplification bias associated with low starting material. It was therefore excluded from downstream analysis. For transparency, expression values for all samples, including B3, are provided in **Supplementary Table S4**. Gene-level abundance was quantified using featureCounts (Subread package v2.0.1), summarizing paired-end reads at the exon level based on Ensembl gene annotations. Differential gene expression analysis was performed between trabecular and compact cardiomyocytes (CMs) using the DESeq2 package (v1.38.0). Genes with less than 10 reads across analyzed samples were filtered out. Differential expressions were calculated between trabecular CMs (Double RFP+/GFP+) and compact CMs (RFP+) using the Wald test. Subsequently, p-values were adjusted for multiple testing using the Benjamini-Hochberg method. Genes were considered significantly differentially expressed at an adjusted p-value (padj) < 0.05. Gene Ontology (GO) enrichment analysis was performed on significantly up-regulated and down-regulated genes in trabecular CMs using the clusterProfiler package in R. Further, p-values were adjusted using the Benjamini-Hochberg method and the terms with an adjusted p-value (padj) < 0.05 were considered significant.

#### Building of protein-protein interaction and its analysis

Protein-Protein Interaction Network (PPIN) was constructed using the STRING database application (v11.5) within Cytoscape (v3.10.4) using the list of significantly differentially expressed genes (padj < 0.05) as the query input. Nodes in the resulting PPIN represent encoded proteins and edges denote functional associations, including both physical interactions and functional relationships derived from experimental evidence, co-expression and text-mining. Edge thickness represents confidence score, with thicker lines indicating stronger evidence of interaction and vice versa. Nodes are coloured based on their log2FC values such as down-regulated genes (enriched in compact CMs) are shown in a gradient from dark to light green, up-regulated genes (enriched in trabecular CMs) in a gradient from dark to light red and genes with nominal fold changes (log2FC ± 0.12) are coloured white. Next, PPIN was clustered to obtain tightly interacting protein modules using MCODE algorithm.

### Statistical analysis

Normality was assessed using the Shapiro–Wilk test and homogeneity of variances using the F-test (for two groups) or Levene’s test (for multiple groups). Two-group comparisons were performed with Student’s *t*-test; Welch’s *t*-test was used when variances were unequal, and the Wilcoxon rank-sum test was applied when normality was not assumed. For comparisons across more than two groups, one-way ANOVA followed by Tukey’s HSD test was used. Fisher’s exact test was applied to compare EGFP+ distributions between conditions. Statistical analyses were performed in R. All statistical analyses were performed using the R statistical programming language. All the plots were generated using RStudio software^17^ and Microsoft Excel.

## Data availability

All sequencing data generated in this study have been deposited in the GEO database under accession number GSE315841. Processed data are provided as Supplementary Data.

## Acknowledgements

We are grateful to the Zebrafish Core Facility of the IIMCB Warsaw for excellent fish care and Microscopy and Cytometry Facility for experimental support; V. Korzh, J. Kuznicki, P. Gawlinski, J. Wysocka, and D. Stainier, as well as all members of the Winata lab for fruitful discussions and critical guidelines.

## Author Contributions

C.P. performed experiments and analyzed data. S.V., K.G.S., and K.M. performed transcriptomics analysis. M.Z. and M.G.N. performed spatial analysis. C.W., C.P., and M.Z. wrote the manuscript. C.W. conceived and supervised the project and designed the experiments. All authors have read and approved the final paper.

## Additional Information Funding

This research was supported by National Science Center Poland, grant number 2018/29/B/NZ2/01010L and 2019/35/B/NZ2/02548. M.S.G.N and M. Z. were supported by a grant from the Priority Research Area DigiWorld under the Strategic Program Excellence Initiative at Jagiellonian University and by the National Science Center Poland, grant number 2021/42/E/NZ2/00188. All transgenic zebrafish lines used in this study were maintained at the IIMCB zebrafish core facility (IN-MOL-CELL Infrastructure) funded by the European Union – NextGenerationEU under National Recovery and Resilience Plan. IN-MOL-CELL Infrastructure was also funded by the European Union under Horizon Europe (project 101059801 - RACE) and RACE-PRIME project carried out within the IRAP programme of the

Foundation for Polish Science co-financed by the European Union under the European Funds for Smart Economy 2021–2027 (FENG).

