## Supplemental Materials for "Enhancer-mediated metabolic pre-patterning defines trabecular cardiomyocyte identity prior to morphogenesis"

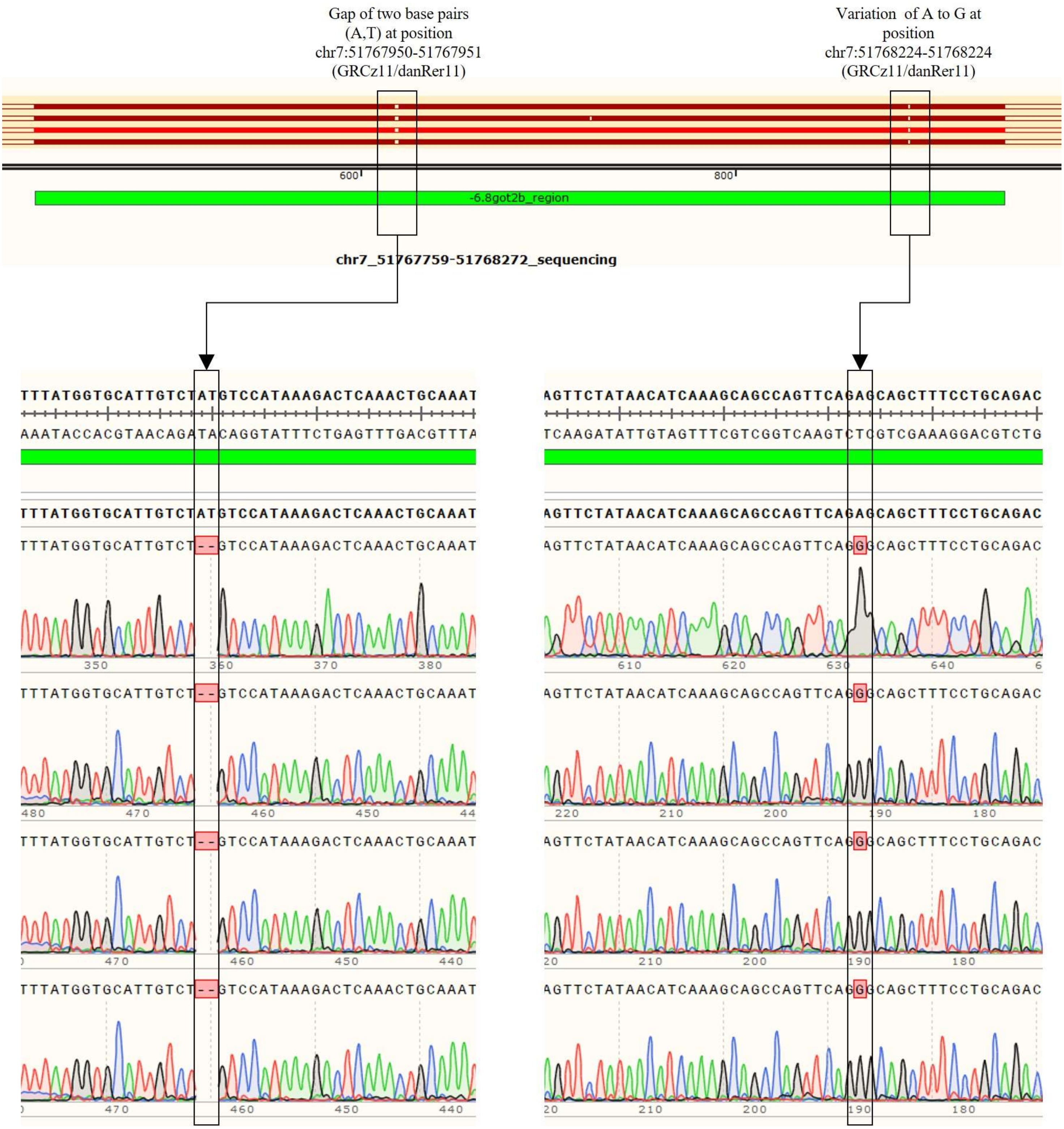


Supplemental figure S1

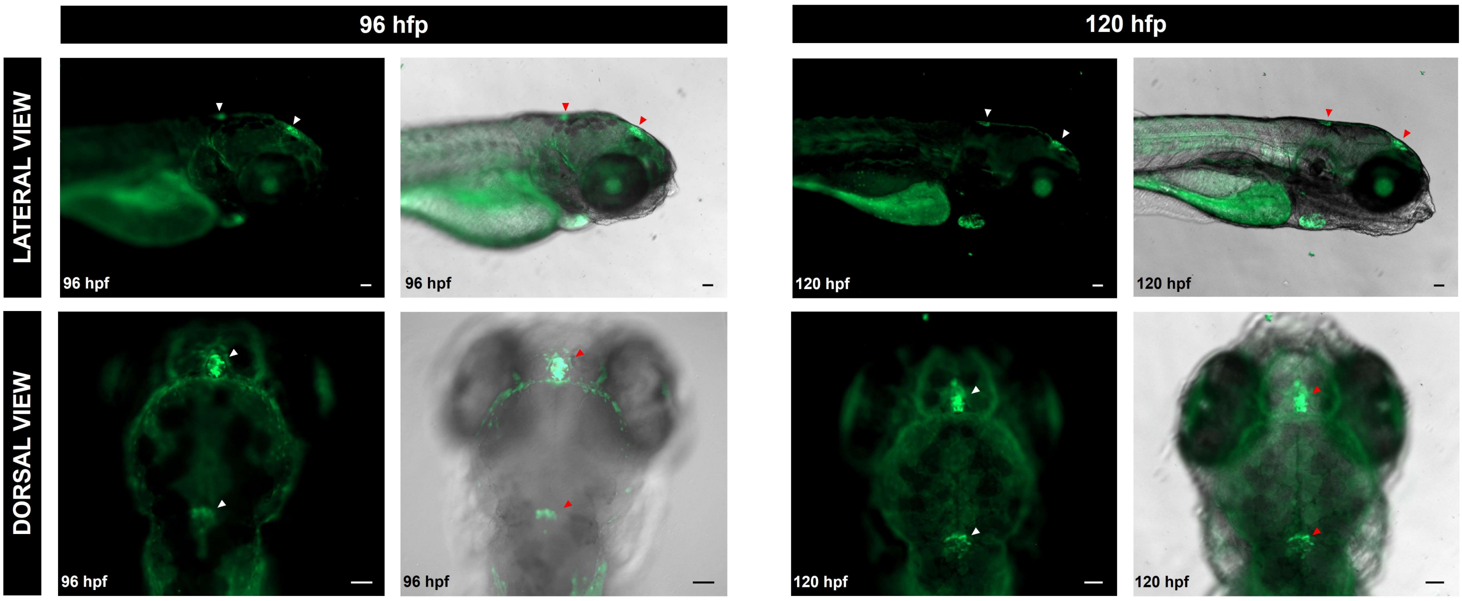


Supplemental figure S2

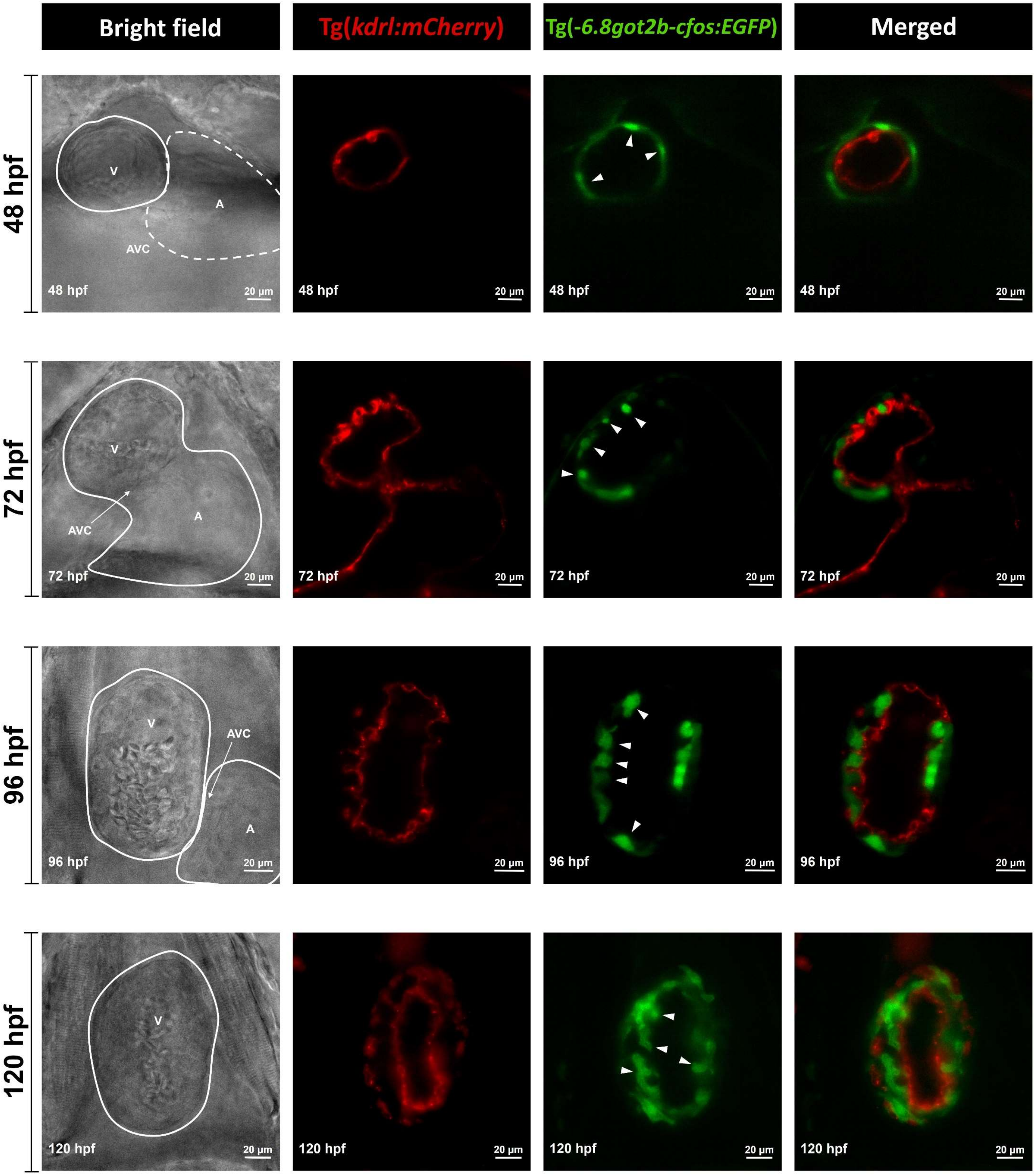


Supplemental figure S3

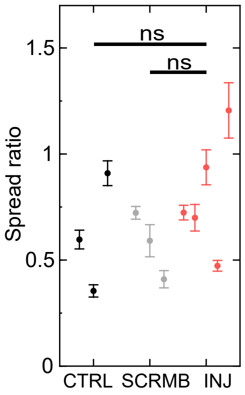
ns
ns
1.5
1
0.5
0
Spread ratio
CTRL
SCRMB
INJ

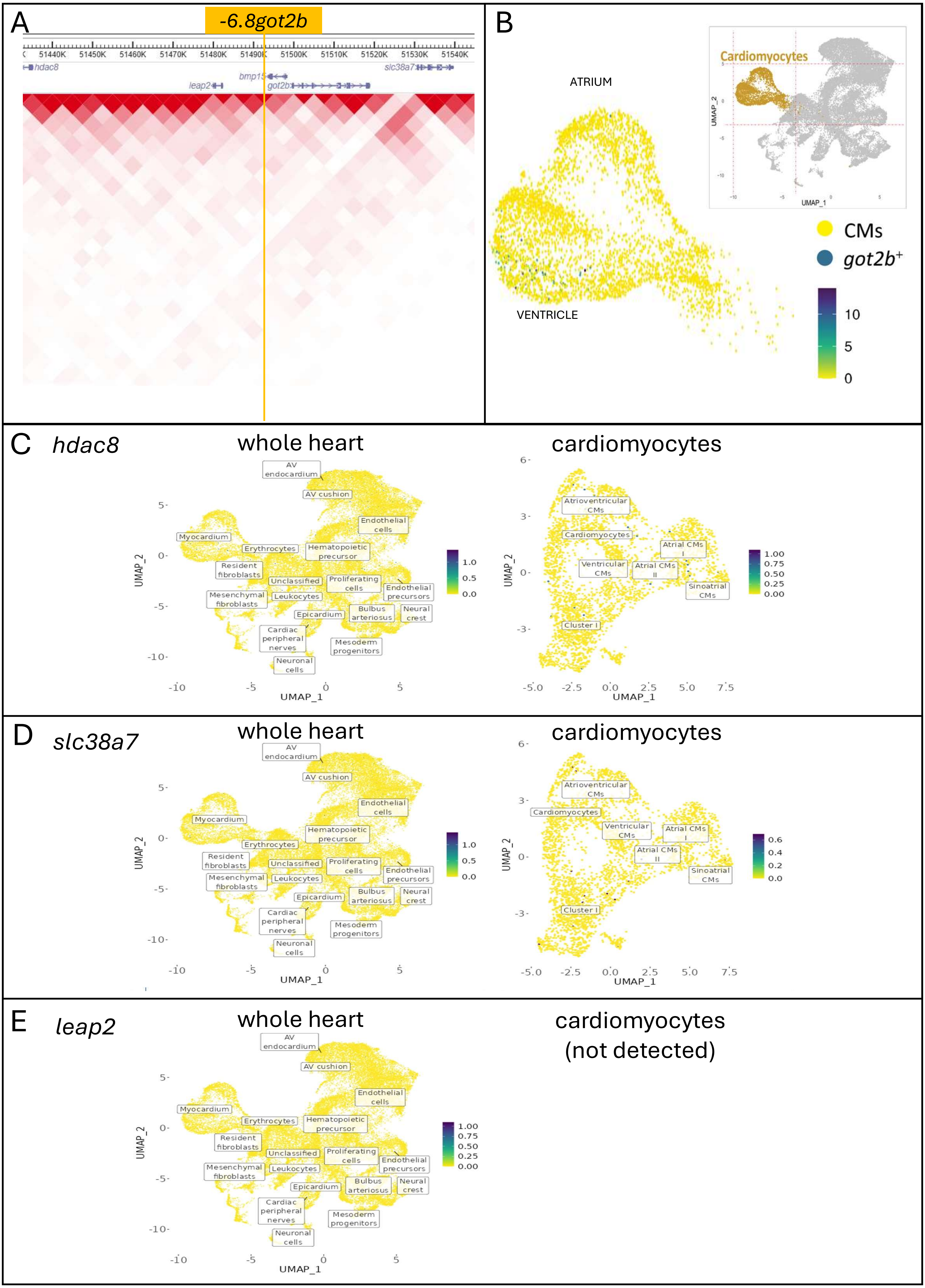


Supplemental figure S5

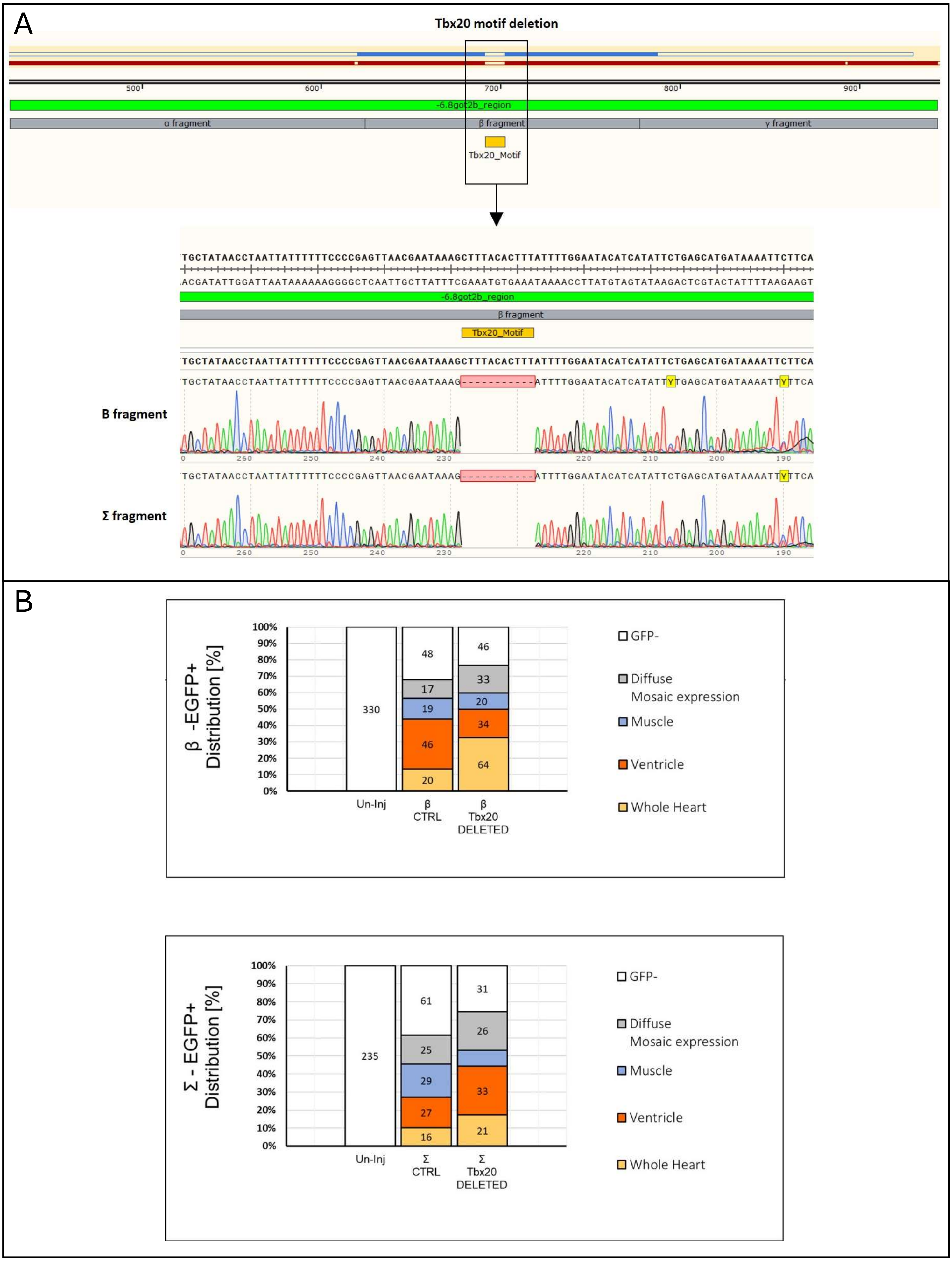


Supplemental figure S6

A

B

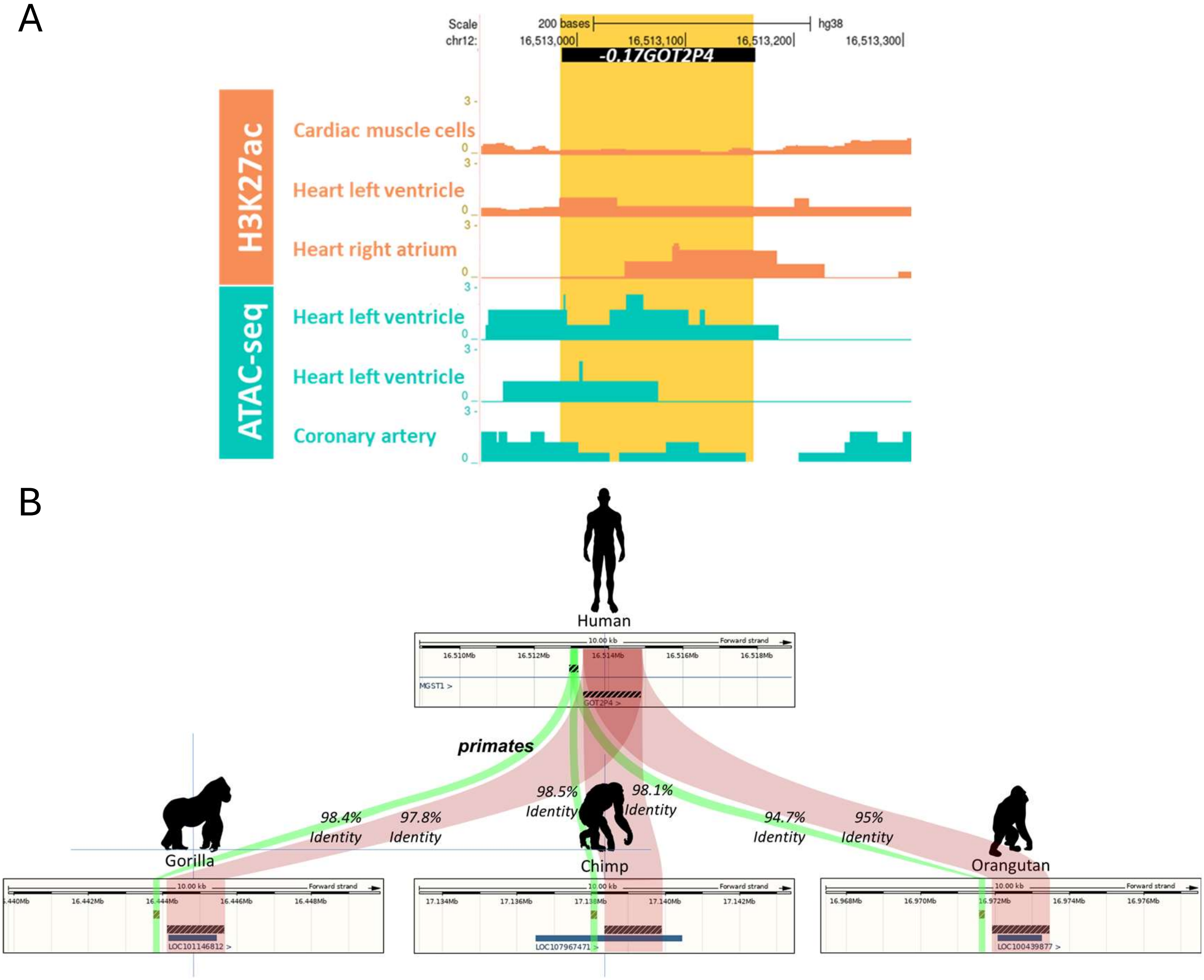


Supplemental figure S7

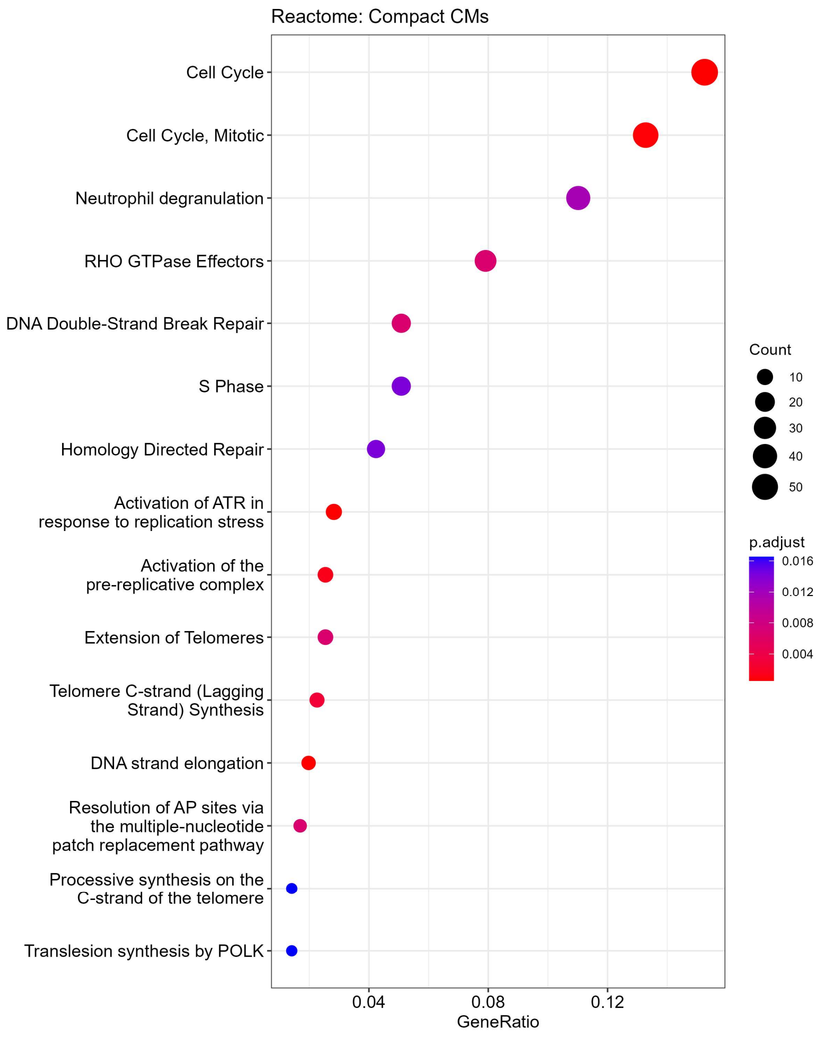


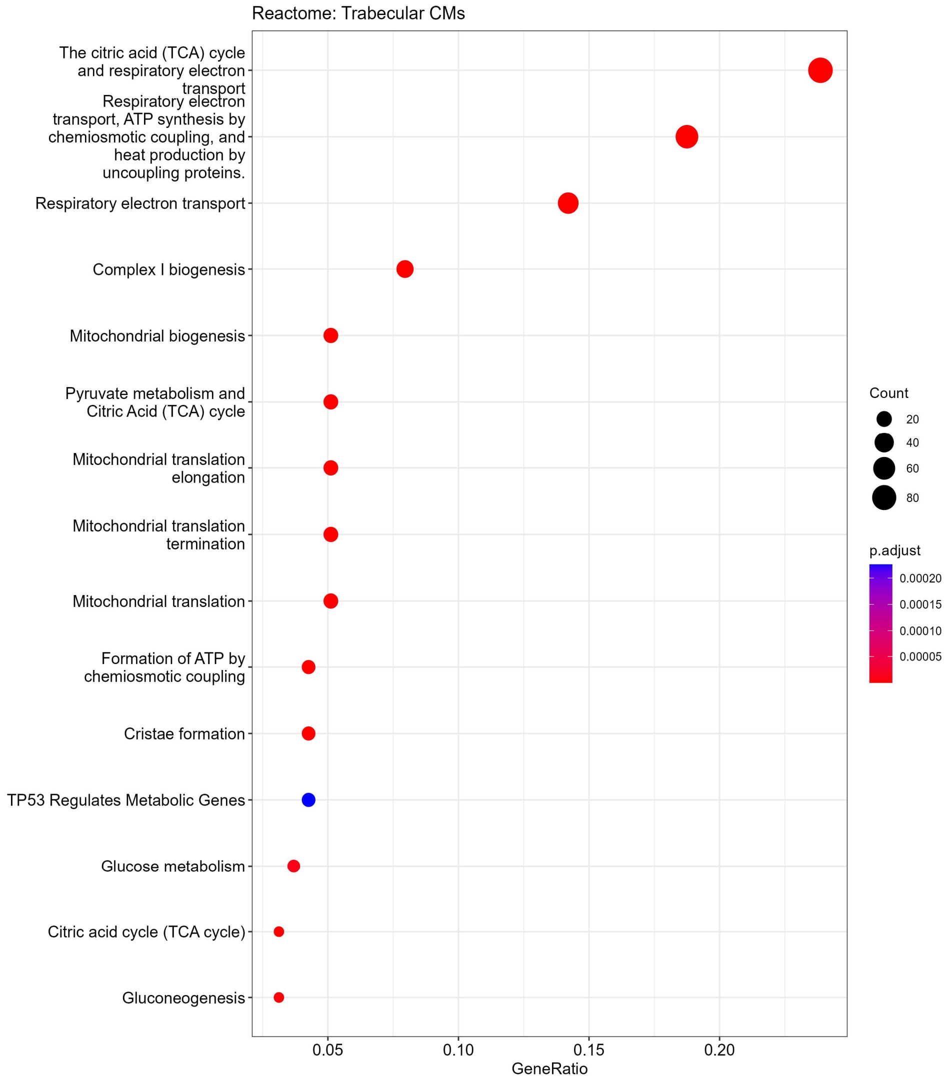


### Supplemental figure S8

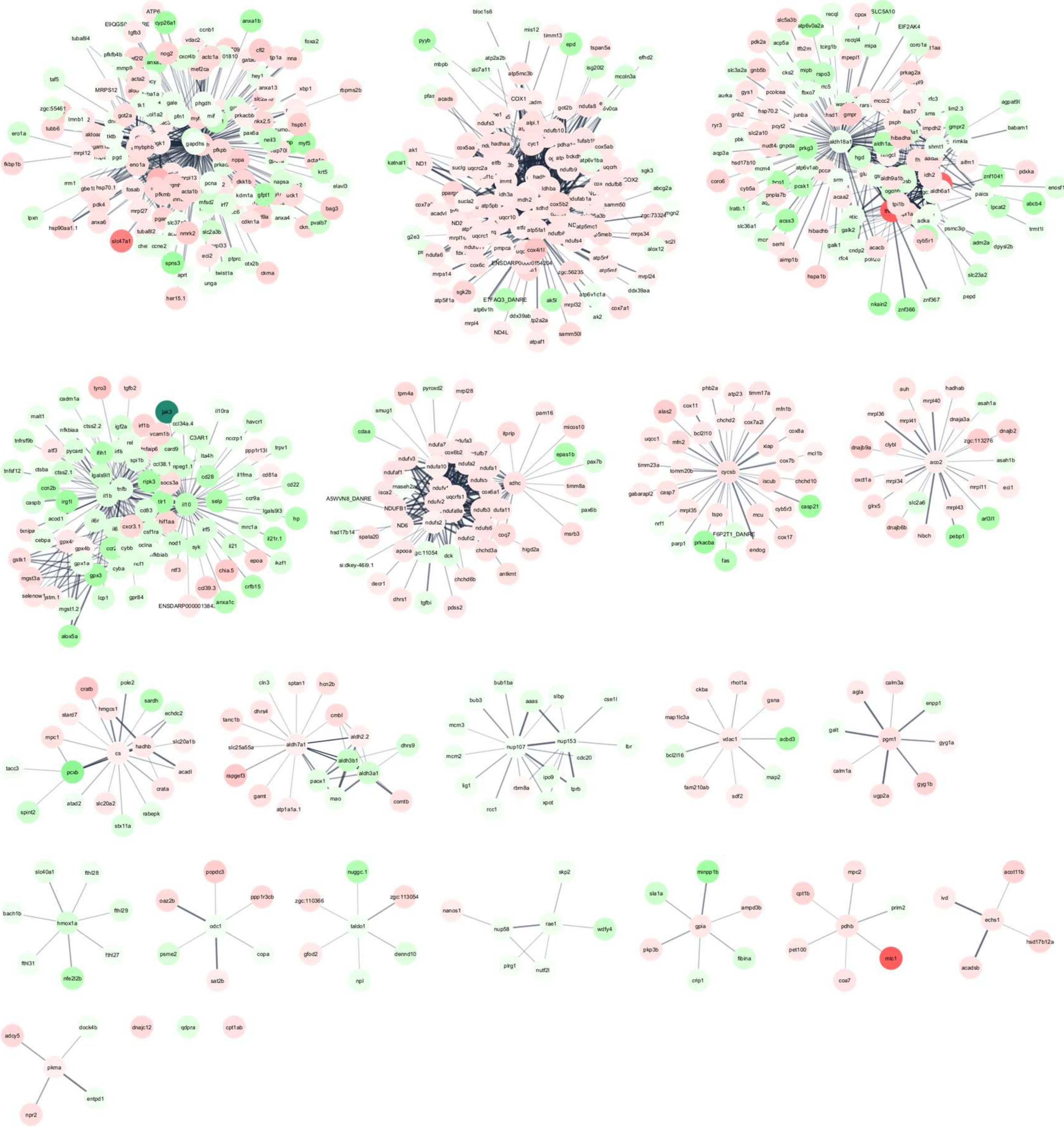


Supplemental figure S9
